# Evolution guided tolerance engineering of *Pseudomonas putida* KT2440 for production of the sustainable aviation fuel precursor isoprenol

**DOI:** 10.1101/2025.02.28.640801

**Authors:** Hyun Gyu Lim, Aparajitha Srinivasan, Russel Menchavez, Ian S. Yunus, Myung Hyun Noh, Megan White, Yan Chen, Jennifer W. Gin, Bernhard O. Palsson, Taek Soon Lee, Christopher J. Petzold, Thomas Eng, Aindrila Mukhopadhyay, Adam M. Feist

**Affiliations:** Department of Bioengineering, University of California San Diego, 9500 Gilman Dr., La Jolla, CA 92093, USA; Joint BioEnergy Institute, 5885 Hollis street, 4th floor, Emeryville, CA 94608, USA; Department of Biological Sciences and Bioengineering, Inha University, Inha-ro 100, Michuhol-gu, Incheon 22212, Korea; Biological Systems and Engineering Division, Lawrence Berkeley National Laboratory, 1 Cyclotron Road, Berkeley, CA 94702, USA; Novo Nordisk Foundation Center for Biosustainability, Technical University of Denmark, 2800 Kgs, Lyngby, Denmark; Environmental Genomics and Systems Biology Division, Lawrence Berkeley National Laboratory, 1 Cyclotron Road, Berkeley, CA 94702, USA

**Keywords:** Adaptive laboratory evolution, *Pseudomonas putida*, isoprenol, 3-methyl-3-buten-1-ol, sustainable aviation fuel (SAF), tolerance engineering

## Abstract

Isoprenol (3-methyl-3-buten-1-ol) is a sustainable aviation fuel precursor and commodity chemical which can be microbially synthesized from renewable carbon streams. Its production has been demonstrated in *Pseudomonas putida* KT2440 but its titers, rates, and yields have yet to reach commercially viable levels, potentially due to its inherent toxicity. We hypothesized that utilization of Tolerization Adaptive Laboratory Evolution (TALE) would generate *P. putida* hosts more tolerant to isoprenol and suitable for enhanced production phenotypes. Here, we performed a comprehensive TALE campaign using three strains, the wild-type and two strains lacking subsets of known isoprenol catabolism and transport functions in quadruplicate independently evolved lineages. Several evolved clones from each starting strain displayed robust growth (up to 0.2 h^−1^) at 8 g/L of isoprenol, where starting strains could not grow. Whole genome resequencing of the 12 independent strain lineages identified convergent mutations. Reverse engineering four commonly mutated regions individually (*gnuR*, *ttgB*-PP_1394, PP_3024-PP_5558, PP_1695) each resulted in a partial recovery of the tolerance phenotypes observed in the evolved strains. Additionally, a proteomics-guided deletion of the master motility regulator, *fleQ*, in an evolved clone alleviated the tolerance vs. production trade-off, restoring isoprenol titers and consumption to levels observed in the starting strains. Collectively, this work demonstrated that an integrated strategy of laboratory evolution and rational engineering was effective to develop robust biofuel production hosts with minimized product toxicity.

## 1. Introduction

Isoprenol (3-methyl-3-buten-1-ol) is an established advanced biofuel and a precursor for diverse commodity chemicals (Zheng et al., 2013), and can be produced from biomass using heterologous pathway expression in a microbial host. Chemical catalysis can efficiently convert isoprenol to isoprene, a key component for synthetic rubber synthesis and more broadly, a drop-in platform chemical compatible with existing chemical manufacturing workflows (Emelianov et al., 2024; Kant et al., 2023). As a fuel molecule, it has a higher energy density compared to ethanol (36.5MJ/kg vs 36.8 MJ/kg) (Pang et al., 2021). Moreover, it can be converted into 1,4-dimethylcyclooctane (DMCO), an emerging sustainable aviation fuel (Baral et al., 2021; Benavides et al., 2022; Rosenkoetter et al., 2019). DMCO has a 9.2% higher volumetric net heat of combustion compared to that of the currently used JET-A (Baral et al., 2021; Rosenkoetter et al., 2019). Currently, the wide adoption of isoprenol is limited as existing bioconversion processes lack the high titers, rates, and yields to derisk its production at scale using renewable carbon streams.

*Pseudomonas putida* is a versatile microbial chassis with rapid growth, broad substrate utilization (e.g., coumarate, ferulate, benzoate), and high-stress tolerance, making it ideal for bioprocesses using biomass-derived carbon streams (Belda et al., 2016; Nikel and de Lorenzo, 2018; Wada et al., 2021). Targeted host engineering of *P. putida* has allowed efficient utilization of both five and six carbon sugars including xylose, arabinose, and galactose (Elmore et al., 2020; Lim et al., 2021; Mohamed et al., 2020). Furthermore, a range of systems biology approaches have enabled a better understanding of its transcriptional regulatory network (Borchert et al., 2024; Lim et al., 2022) informing new strain designs with sophisticated tools for genetic engineering (Martin-Pascual et al., 2021; Nikel et al., 2014). Currently, *P. putida* has been demonstrated as a viable host for a range of biochemicals including PHA (Cha et al., 2020), lycopene (Hernandez-Arranz et al., 2019), indigoidine (Banerjee et al., 2020; Eng et al., 2023; Gauttam et al., 2023; Lim et al., 2021), muconic acid (Almqvist et al., 2021; Bentley et al., 2020; Ling et al., 2022; van Duuren et al., 2011), lactic acid (Zou et al., 2021), adipic acid (Ackermann et al., 2021; Niu et al., 2020), medium-chain fatty acid methyl esters (Valencia et al., 2022) and more recently, isoprenol (Banerjee et al., 2024; Wang et al., 2022).

One critical challenge in using *P. putida* as an isoprenol production platform is product toxicity, despite its general tolerance to other commodity chemicals. Although it is known to exhibit relatively higher tolerance to hydrophobic compounds than model bacteria including *Escherichia coli*, cell growth is still inhibited since alcohols including isoprenol denature proteins and increase membrane instability (Huffer et al., 2011). Tolerizing microbes for high concentrations of a shorter alcohol, ethanol, has resulted in the identification of both degradative routes (*i.e.* increased or new enzymes capable of catabolism) and non-degradative routes (*i.e.*, membrane fluidity, cell wall modifications, induction of stress response systems, chaperones, efflux pumps and production of compatible solutes such as glycine-betaine) (Mukhopadhyay, 2015). While previous reports for improved heterologous isoprenol production in *P. putida* exceed 1 g/L (Banerjee et al., 2024), product toxicity may limit further increases as intracellular isoprenol concentrations may approach growth inhibitory levels.

To improve tolerance phenotypes of microbial hosts, Adaptive Laboratory Evolution (ALE) has been widely applied (Sandberg et al., 2017). After continuous exposure to an increasing target inhibitory compound (a specific methodology referred to as Tolerization ALE or TALE) isolates with improved tolerance can be obtained. In addition, subsequent -omics investigations (e.g., genome, transcriptome, or proteome analysis) can suggest tolerization mechanisms, expanding our understanding about microbial physiology. Accordingly, there are many studies which have tolerized microorganisms against diverse growth inhibitors including alcohols (Halle et al., 2023), ionic liquids (Lim et al., 2020; Mohamed et al., 2017), and organic acids (Lennen et al., 2023; Nguyen-Vo et al., 2019). A recent evolution study with *E. coli* K-12 MG1655 has successfully increased the half-maximal inhibitory level against isoprenol by 47% (Babel and Krömer, 2020) and identified tolerance mechanisms with a subsequent mutation analysis. Further, *P. putida* can inherently catabolize isoprenol (Thompson et al., 2020), which could be one of the major tolerance mechanisms. To develop *P. putida* strains suitable for high isoprenol production, it is therefore likely important to explore tolerance mechanisms other than catabolism.

In this study, we performed an in-depth comparative TALE experiment to improve and understand the mechanisms of tolerance of *P. putida* KT2440 (hereafter, KT2440) to isoprenol. In addition to the wildtype KT2440 strain (WT), we constructed two additional starting strains lacking isoprenol catabolism and a putative isoprenol efflux pump to generate tolerized strains via the known routes. All three strains were tolerized to gradually increasing concentrations of isoprenol. We successfully obtained evolved isolates that can robustly grow at 8 g/L isoprenol, a lethal level to WT (despite its ability to catabolize isoprenol) and characterized the resulting system-level changes by DNA resequencing and shotgun proteomics. While isoprenol production was initially low in evolved clones, rational engineering approaches based on proteomics analysis restored isoprenol titers to comparable levels to the starting control strain.

## 2. Materials and Methods

### 2.1 Bacterial cells, plasmids, and reagents

Bacterial strains, plasmids and oligonucleotides used in this study were listed in **Supplementary Table 1** and **2.** Oligonucleotides were synthesized by Integrated DNA Technologies (IDT, Coralville, IA, USA). Plasmid cloning was performed by using a NEBuilder^R^ HiFi DNA Assembly Master Mix from New England Biolabs (NEB, Ipswich, MA, USA). Q5 and HotStartTaq polymerases were also purchased from NEB. Scar-less gene deletion was performed by following the conjugation protocol using *E. coli* S-17 as previously described (Lim et al., 2020). Alternatively, we used recombineering to generate deletion mutants with oligonucleotides in conjunction with a Cpf1/RecT two-plasmid system (Czajka et al., 2022; Wannier et al., 2020). All chemicals were purchased from Sigma-Aldrich (St. Louis, MO, USA) unless otherwise mentioned. We note that commercial isoprenol stock (97%) contains formaldehyde as an impurity (Babel and Krömer, 2020; Dyga et al., 2021).

### 2.2 Bacterial cultivation

All bacterial strains were routinely cultivated on LB medium or the modified M9 minimal medium. The M9 medium contained 2 g/L (NH_4_)_2_SO_4_, 6.8 g/L Na_2_HPO_4_, 3 g/L KH_2_PO_4_, 0.5 g/L NaCl, 2 mM MgSO_4_, 0.1 mM CaCl_2_, 500 μL/L 2000× trace element solution (Lim et al., 2020; Linger et al., 2014). As a carbon source, 4 g/L glucose was added to the minimal medium unless otherwise stated. To monitor cell growth, colonies grown in LB agar plates were picked and inoculated into 15 mL of the M9 medium contained in 30 mL cylindrical flasks. After overnight growth, cultures were re-inoculated at an OD_600_ (optical density at 600 nm) of 0.05. Once OD_600_ reached 0.6-0.8, cells were further inoculated at OD_600_ of 0.05 as main cultures. For small scale cultures (i.e., microtiter plates), subsequent cultures were inoculated at OD_600_ of 0.1 in 200 μL of a culture medium. Maximum growth rates were calculated by plotting slope of ln (OD_600_) vs. time during exponential growth.

TALE was performed by passaging bacterial cultures to 15 mL of fresh medium in flasks once or twice a day at an initial OD_600_ value of 0.05. Cell growth was monitored by measuring OD_600_ of cultures over time. Concentrations of isoprenol were increased from 4 g/L (46 mM) by a step size of 0.5 or 1 g/L. When an increase in isoprenol concentration resulted in complete growth inhibition, the previous culture was passaged in non-selective M9 medium and the experiment was restarted with the prior isoprenol concentration. Each evolved isolate was named according to the ALEdb convention (Phaneuf et al., 2019) that uses the ALE experiment number (A), flask number (F), and isolate number (I).

### 2.4 Genome and transcriptome sequencing

For whole-genome sequencing, genomic DNA was extracted from 500 μL of saturated cell cultures by using a Mag-Bind^R^ Bacterial DNA 96 Kit from Omega Bio-tek (Norcross, GA, USA) and KingFisher™ Duo Prime Purification System from Thermo Fisher Scientific (Waltham, MA, USA). Genomic sequencing libraries were prepared by using a plexWell™ 96 kit from seqWell (Beverly, MA, USA). Raw files were processed by an in-lab analysis pipeline that utilizes Breseq (version 0.33.1) (Deatherage and Barrick, 2014), GATK (Van der Auwera and O’Connor, 2020; Van der Auwera et al., 2013), and Bowtie2 (Langmead and Salzberg, 2012) (version 2.3.4.1). Mutation analysis results were uploaded to ALEdb v1.0 (Phaneuf et al., 2019) and accessible at http://aledb.org/ with “Pputida_isoprenol_TALE” as the project name. Sequencing coverage varied between 10 - 50X; the coverages of endpoint clones were at least 30X. Starting strain mutations were filtered out. Raw-read files were deposited to NCBI SRA with a BioProject number of PRJNA1187681.

For transcriptome sequencing, the log phase WT cells grown on either glucose or isoprenol as a sole carbon source were taken. The total RNA was extracted from 3 mL of cell cultures at the exponential growth phase (OD_600_ 0.6-0.8), stabilized by mixing 6 mL of an RNAprotect Bacteria Reagent from Qiagen (Hilden, Germany). Total RNA was extracted by using a Quick-RNA Fungal/Bacterial Microprep Kit from Zymo (Irvine, CA). Subsequently, ribosomal RNA was depleted by following the RiboRid (Choe et al., 2021) and OligoRid protocol using anti-ribosomal RNA oligonucleotides designed for KT2440 (Lim et al., 2020). Transcriptome sequencing libraries were prepared by using either a Swift Rapid RNA Library Kit from Swift (Ann Arbor, MI) or KAPA RNA HyperPrep Kit from Roche Sequencing (Pleasanton, CA). Prepared libraries were sequenced by using an illumina (San Diego, CA) NextSeq or NovaSeq platform. Raw reads were analyzed by using Bowtie2, DESeq2 (Love et al., 2014), summarizeOverlaps (Lawrence et al., 2013). Raw read files were deposited at Gene Expression Omnibus with an accession number of GSE281392.

### 2.5 Biomass quantification

Biomass was determined by monitoring absorbance at 600 nm (OD_600_) by using an Infinite 200 PRO microtiter plate reader from Tecan (Männedorf, Switzerland), BioTek Synergy microplate reader (Agilent Technologies, USA), or a Biomate 3S benchtop spectrophotometer from Thermo Fisher Scientific (Waltham, MA, USA) wherever appropriate. Microtiter plate OD_600_ readings measured by the Infinite 200 Pro microtiter plate reader were multiplied by a factor of 0.2211 for conversion into readings by the bench-top spectrophotometer.

### 2.6 Shotgun proteomic analysis

Cultures at the exponential growth phase were harvested by centrifugation and stored at −80 °C until analysis. After all samples were collected, protein was extracted from the pellets and tryptic peptides were prepared by following established proteomic sample preparation procedures (Chen et al., 2023). Briefly, cell pellets were resuspended in Qiagen P2 Lysis Buffer (Qiagen Sciences, Germantown, MD, Cat.#19052) for cell lysis. Proteins were precipitated with the addition of 1 mM NaCl and 4 x volume acetone, followed by two additional washes with 80% acetone in water. The recovered protein pellet was homogenized by pipetting mixing with 100 mM ammonium bicarbonate in 20% methanol. Protein concentration was determined by the DC protein assay (Bio-Rad Inc, Hercules, CA). Protein reduction was accomplished using 5 mM Tris-2-(carboxyethyl) phosphine for 30 min at room temperature, and alkylation was performed with 10 mM iodoacetamide as a final concentration for 30 min at room temperature in the dark. Overnight digestion with trypsin was accomplished with a 1:50 trypsin:total protein ratio. The resulting peptide samples were analyzed on an Agilent 1290 UHPLC system coupled to a Thermo Scientific Orbitrap Exploris 480 mass spectrometer for discovery proteomics (Chen et al., 2022). Briefly, peptides were loaded onto an Ascentis ES-C18 Column (Sigma–Aldrich, St. Louis, MO) and separated with a 10 min LC gradient starting at 98% solvent A (0.1 % FA in H2O) and 2% solvent B (0.1% FA in ACN) to 65% solvent A and 35% solvent B. Eluting peptides were introduced to the mass spectrometer operating in positive-ion mode and were measured in data-independent acquisition (DIA) mode with a duty cycle of 3 survey scans from m/z 380 to m/z 985 and 45 MS2 scans with precursor isolation width of 13.5 m/z to cover the mass range. DIA raw data files were analyzed by an integrated software suite DIA-NN. The database used in the DIA-NN search (library-free mode) is the latest Uniprot *P. putida* KT2440 proteome FASTA sequence plus the protein sequences of heterogeneous pathway genes and common proteomic contaminants. DIA-NN determines mass tolerances automatically based on first pass analysis of the samples with automated determination of optimal mass accuracies. The retention time extraction window was determined individually for all MS runs analyzed via the automated optimization procedure implemented in DIA-NN. Protein inference was enabled, and the quantification strategy was set to Robust LC = High Accuracy. Output main DIA-NN reports were filtered with a global FDR = 0.01 on both the precursor level and protein group level. The Top3 method, which is the average MS signal response of the three most intense tryptic peptides of each identified protein, was used to plot the quantity of the targeted proteins in the samples (Ahrné et al., 2013; Silva et al., 2006). With this method, an average of 2,375 out of 5,565 putative proteins were quantified.

The generated mass spectrometry proteomics data have been deposited to the ProteomeXchange Consortium via the PRIDE partner repository with the dataset identifier PXD054609 (Perez-Riverol et al., 2022). DIA-NN is freely available for download from https://github.com/vdemichev/DiaNN.

### 2.7 Isoprenol production and metabolite quantification

For isoprenol production, *P. putida* strains were transformed with approximately 100ng purified plasmid pIY670 (Banerjee et al., 2024) via electroporation (Iwasaki et al., 1994) and selected on solid LB agar containing 50 µg/mL kanamycin. Plasmid pIY670 was prepared for electroporation by extraction from *E. coli* XL-1 using standard alkaline lysis and silica column purification (Qiagen Inc, Plasmid DNA Miniprep Kit). After transformation, a single well-formed *P. putida* colony was picked from the agar plate and used to inoculate a 5 mL LB culture in a test tube which was continuously shaken at 200 rpm and maintained at 30 °C. Cultures were adapted twice in NREL M9 minimal medium (Linger et al., 2014) by back dilution roughly every 24 hours, aiming for robust cell growth. At the third back-dilution, cells were inoculated to a starting OD_600_ of 0.1 in 20-mL test tubes containing 5 mL NREL M9 minimal medium as described in **Section 2.2** with 20 g/L glucose as carbon source and 50 µg/mL kanamycin. The isoprenol pathway was induced by adding arabinose at 2 g/L at T= 0 h. 200 µL of the culture was harvested at 12 h, 24 h and 48 h for analysis of isoprenol, residual glucose and proteomics respectively. Extraction and analysis of isoprenol was carried out as described in a previous study (Banerjee et al., 2024). Briefly, whole cells were extracted with equal volumes of ethyl acetate, vortexed for 15 min and centrifuged at 14,000 *✕ g* for 3 min. 100 µL of the upper organic phase was analyzed using gas chromatography connected to a flame ionization detector (GC-FID) with the following settings - column: 15 m x 0.32 mm x 0.5 µm polar DB-Wax column (Agilent Technologies, USA), carrier gas: helium at a flow rate of 2.22 mL/min; flame: hydrogen; oven temperature gradient: initial hold at 40 °C for 1 min, ramped up at the rate of 15 °C/min to 100°C and finally heated to 230 °C and held at this temperature for 3 min. Isoprenol was quantified from a standard calibration curve using an authentic standard (Sigma Aldrich, USA). Residual glucose and organic acids (lactate, acetate and succinate) were measured using an Agilent 1260 HPLC (Agilent Technologies, USA) equipped with an Aminex HPX-87H column with 4 mM sulfuric acid as the mobile phase at 0.6 mL min^−1^ and analyzed using a refractive index detector (RID). Standard calibration curves were generated using serial dilutions of the stocks of the respective authentic analytes. Samples were analyzed as three independent biological replicates.

## 3. Results and Discussion

### 3.1 Design of an Isoprenol TALE Campaign with Different Starting Strains based on RB-TnSeq and Transcriptome Analysis

RB-Tn-Seq and RNA-Seq analyses were utilized to identify pathways and transporters related to isoprenol metabolism in *P. putida* and to generate multiple starting strains for the TALE campaign (**Figure 1A-C**). We hypothesized that KT2440 employed two potential mechanisms to mitigate isoprenol toxicity: (1) utilization of isoprenol as a carbon source through catabolism, and (2) secretion of isoprenol from the cytoplasm using a transporter, based on protein homology with a probable isoprenol efflux pump, T1E-0242 (TtgB), from *P. putida* DOT-T1E (Basler et al., 2018). We designed gene deletion mutants using known tolerance mechanisms from the literature and complemented them with additional RNA-Seq analysis in response to exogenous isoprenol. We identified differential gene expression targets that were in good agreement with previous RB-TnSeq fitness profiling data (Thompson et al., 2020) (**Supplementary Figure 1A**). From this downselection, we inactivated the pyrroloquinoline quinone (PQQ) cofactor regeneration enzyme PP_2675, as previously reported (Thompson et al., 2020), which limits most alcohol dehydrogenase activity and blocks growth on isoprenol. This approach contrasts with knocking out the approximately 20 promiscuous alcohol dehydrogenases according to UniProt and KEGG annotations. We also deleted genes encoding PP_4064-PP_4067 *ivd*-*mccB*-*liuC*-*mccA* (isovaleryl-CoA dehydrogenase, methylcrotonyl-CoA carboxylase β, methylcrotonyl-CoA hydratase, methylcrotonyl-CoA carboxylase subunit α, respectively) as these encoded proteins may enable the stepwise degradation of isoprenol into acetyl-CoA (**Figure 1B**) (Thompson et al., 2020). Furthermore, the *ttgB* homolog (PP_1385) from DOT-T1E was identified using a simple BLASTn query and deleted in IPL400. Finally, we included PP_3839 as a gene deletion target since it has known activity on other short-chain alcohols (Fong et al., 2006), even though it alone has no fitness defect on isoprenol (**Supplementary Figure 1B**). This led us to construct two stacked deletion strains (IPL300, IPL400) with varying degrees of inactivated isoprenol-related activities compared to the WT (**Materials and Methods, Figure 1B-E**). IPL300 contains deletions in ΔPP_2675, ΔPP_4064-4067, and ΔPP_3839 while IPL400 also contains the same ΔPP_2675, ΔPP_4064-4067, and ΔPP_3839 deletions as well as an additional deletion in Δ*ttgB*.

**Figure 1.**
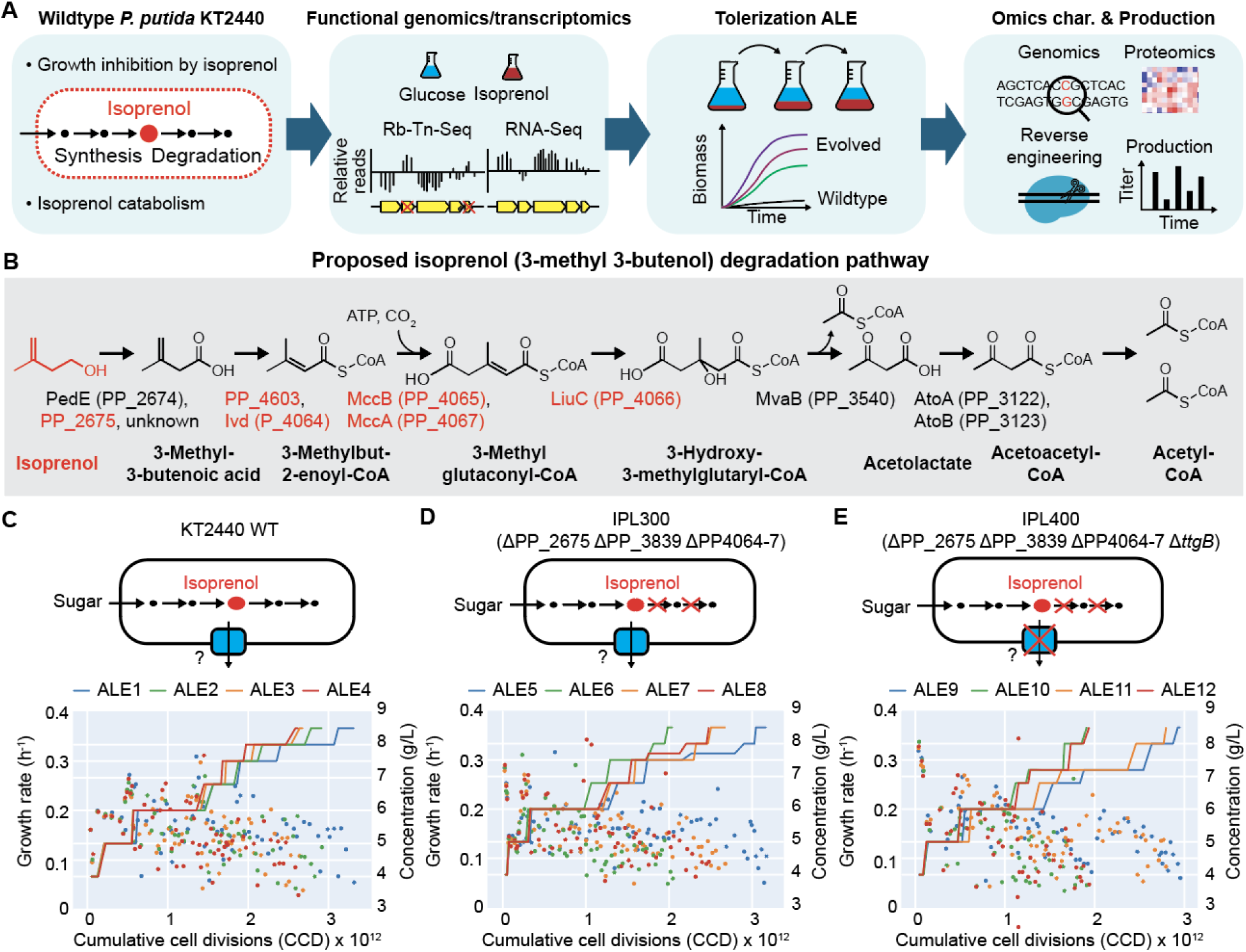
The workflow and rationale for strain selection for TALE to enhance tolerance to isoprenol in *P. putida.* (**A**) Overall workflow for generating and evaluating isoprenol tolerant strains. (**B**) The proposed isoprenol catabolism pathway in *P. putida* (Thompson et al., 2020). Genes targeted for deletion to build the starting strains for TALE are denoted in red. (**C-E**) TALE trajectories of (**C**) KT2440 WT, (**D**) IPL300, (**E**) IPL400. *x*-axes indicate cumulative cell divisions (CCD). Left and right y-axes indicate growth rates, points (h^−1^) of evolving populations and isoprenol concentration, solid lines (g/L), respectively. Growth rates were calculated as described in section 2.5 and had high variability due to limited sampling during the evolutions (Materials and Methods).

The WT and stacked deletion strains IPL300 and IPL400 were phenotypically characterized to understand their tolerance towards exogenously supplied isoprenol in batch culture (**Supplementary Figure 1C-F**). We observed an approximately 2-fold reduction in biomass formation of the WT during the first 16 hours in M9 glucose when it was supplemented with 4 g/L isoprenol (**Supplementary Figure 1C**). This effect became more pronounced with longer lag times at higher isoprenol concentrations (6-10 g/L). Growth completely ceased at 15 g/L. The stacked deletion strains IPL300 and IPL400 showed similar tolerance to isoprenol as the WT. IPL300 exhibited similar growth rate kinetics as the WT in the presence of 4 g/L exogenous isoprenol in glucose minimal medium (**Supplementary Figure 1D**). However, the IPL400 strain, which has an additional *ttgB* deletion, displayed a shortened lag phase and higher final OD_600_ values compared to both IPL300 and WT. This observation was contrary to our expectations and published reports that the strain would be more sensitive to isoprenol due to the lack of this active efflux pump (Basler et al., 2018). Both IPL300 and IPL400 were unable to grow with isoprenol as the sole carbon source in M9 media as they contained deletions of ΔPP_2675 and ΔPP_4064-ΔPP_4067, which were both shown to abolish growth individually on isoprenol when deleted (**Supplementary Figure 1E**). These genes likely encode non-redundant activities given the observation that the ΔPP_2675 strain, but not the ΔPP_4064-ΔPP_4067 strain, essentially blocked isoprenol degradation (**Supplementary Figure 1F**).

### 3.2. Generation of Evolved Strains with Improved Isoprenol Tolerance

We performed high throughput TALE experiments using the WT, IPL300, and IPL400 starting strains in glucose M9 minimal medium by gradually increasing the isoprenol concentration from 4 g/L to 8 g/L in steps of 0.5-1 g/L. Each TALE experiment was conducted with four independent biological replicates in one high-throughput evolution campaign (**Figure 1C** and **Supplementary Table 3;** section 2.3 **Materials and Methods).** The TALE experiments were terminated when the concentration reached 8.0 or 8.5 g/L of isoprenol, as the growth rates were reduced to approximately 0.1 h^−1^ and no further improvements were observed in the tolerized lineages (*i.e.*, the independent biological replicates) after approximately ∼20 generations at the final isoprenol concentrations. We also tolerized the WT strain against formaldehyde stress as a control (HCHO, isolates A13-A16, **Figure 2A** and **2B**) since formaldehyde is a trace impurity in commercially synthesized isoprenol used in this study (Babel and Krömer, 2020; Dyga et al., 2021). Over an approximate 2-month period, these *P. putida* strains underwent approximately 143 to 353 generations, which is equivalent to 1.33 to 3.32 × 10^12^ cumulative cell divisions (CCD). Similarly, for the HCHO TALE, the concentration of HCHO was increased from 0.03 g/L to 0.27 g/L over 346-387 generations, corresponding to 3.11 × 10^12^ to 3.56 × 10^12^ CCDs. At the end of the TALE campaign, end-point clones were isolated from the final flasks of each of the 16 lineages for further phenotype and genotype analysis.

**Figure 2.**
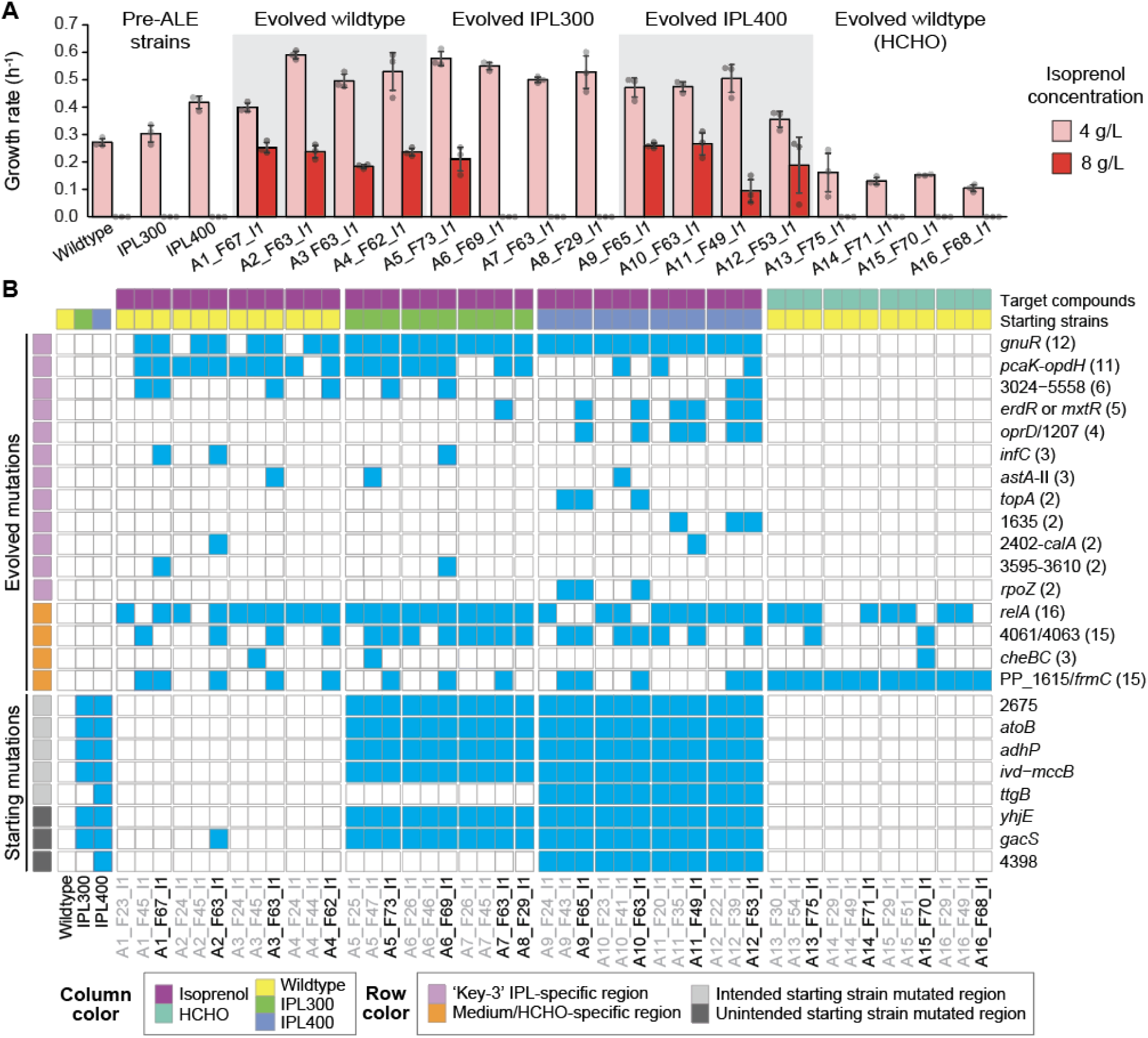
Growth rates of the starting strains and evolved isolates in the presence of isoprenol and mutations identified in isolated clones. (**A**) Maximum specific growth rates (h^−1^) of the starting strains and the evolved endpoint strains in the presence of 4 g/L and 8 g/L of isoprenol in flasks. (**B**) Mutated genes identified in the starting, intermediate (gray font) and endpoint evolved isolates. Refer to Supplemental Table 3 for information on the specific type of mutation(s) recovered in each genetic locus.

To confirm improved fitness of the evolved isolates against their respective tolerized target compound, we compared the growth of the end-point isolates at 4 g/L or 8 g/L exogenous isoprenol with the three starting strains (isolates A1-A12, **Figure 2A**). Notably, all the evolved isolates displayed higher growth rates (0.50 h^−1^, 1.8-fold increase on average) than that of the wild-type strain at 4 g/L isoprenol, the starting concentration. At 8 g/L, all evolved isolates except for three evolved IPL300 isolates (isolates A6-A8, **Figure 2A**) showed growth rates of up to 0.25 h^−1^. Not only were the growth rates improved, but the average lag time was also reduced to essentially zero from 1 h in the presence of 4 g/L isoprenol. There was no strict correlation between the enhancement in the growth rates of the endpoint isolates either at 4 g/L or 8 g/L exogenous isoprenol with the strain backgrounds (isoprenol catabolism present or deficient). Many of the evolved IPL300 end-point isolates did not show growth at 8 g/L (**Figure 2A**) even though they were isolated from the populations which were exposed to essentially the same endpoint isoprenol concentration of 8.5 g/L (refer to **Figure 1C-E**). This phenotypic difference was likely owed to differences in the tolerance assay setup, where in the TALE regime populations were grown with constitutive and increasing concentrations of isoprenol (i.e., from an active growing state). In contrast, in the validation assay, naive cells were directly shifted to the higher 8 g/L isoprenol concentration without exposure to a lower intermediate concentration. The screened isolates tolerized against HCHO did not show improvements in tolerance to isoprenol as no HCHO-tolerant clones were cross tolerant to 8 g/L isoprenol. Overall, the validation of TALE end-point isolates using growth rate and lag time as phenotypic metrics confirmed that the isolated mutants showed up to 1.8-fold improvements for growth rate with greatly decreased lag time in the presence of isoprenol for the WT as well as the isoprenol-catabolism deficient IPL300 and IPL400 starting strains.

### 3.3. Mutation Profiling of the Evolved Isolates to Reveal Mechanisms for Improved Tolerance

A mutational analysis of the evolved TALE strains was performed to link their altered phenotypes to mutated genes. After the isoprenol TALE campaign, *P. putida* mutants showed increased growth rates in the presence of 4 g/L isoprenol and most of them were capable of growing in the presence of 8 g/L, which was growth inhibitory to the starting WT, IPL300 and IPL400 strains of *P. putida*. The phenotypic similarity for isoprenol tolerance across multiple lineages suggested a possibility of correlating common mutations across these 12 biologically independent lineages to identify common determinants of isoprenol tolerance. However, it was also possible that strain-specific mutants existed that enabled growth at the 8 g/L isoprenol condition in some strains, owing to the polytropic impact alcohols may have on cell membrane structure and protein denaturation. We therefore performed whole-genome sequencing of evolved isolates for a total of 46 strains (including the HCHO tolerized isolates), representing the endpoint isolates and several randomly chosen isolates from two intermediate time points per lineage (except for A8 due to a small passage number, **Figure 2B** and **Supplementary Dataset 1,** see **Materials and Methods**). We identified a total of 158 unique mutations in 73 genetic regions (i.e., genes or intergenic regions between two genes, **Supplementary Dataset 1**). The evolved end-point isolates acquired 10 mutations on average. None of the strains accumulated mutations in genes responsible for mutation repair systems such as (*e.g.*, *mutL* or *mutS*) that show such hypermutator populations a growth advantage over non-mutators, a common feature of long term ALE experiments in *E. coli* (Kang et al., 2019) and something that can occur at shorter ALE experiments (LaCroix et al., 2015). We also noted that the starting strains IPL300 and IPL400 also contained pre-existing mutations missing from the reference sequence: PP_4986, *gacS*, *yhjE* in IPL300, and then IPL400 had the same mutations along with a mutation in PP_4398. These mutations were likely generated during the strain construction process and point to the fact that many mutations can arise during such strain engineering campaigns.

An analysis of convergent evolution was performed to understand mutational mechanisms enabling enhanced tolerance phenotypes. A total of 20 regions were mutated in isolates from at least two independent lineages across starting strains and for both isoprenol and HCHO (**Supplementary Data 1**). We filtered out mutations from five regions, *galP*-I/PP_1174 (‘/’ indicates an intergenic region), PP_1325, *relA*, PP_3820/*galU*, PP_4061/PP_4063 due to their frequent occurrence in previous evolution studies with glucose-minimal media (Lim et al., 2020; Mohamed et al., 2020; Phaneuf et al., 2019) and are therefore not specifically mutated in the presence of isoprenol. Additional three regions, PP_1615/*frmAC*, *cheBC*, PP_4671, were mutated in isolates from the HCHO-tolerization experiments (**Figure 2B**), indicating that these regions are likely associated with tolerance mechanisms against the trace impurity HCHO. At the end of this down-selection, we identified 12 loci (*gnuR*, *pcaK*-*opdH*, PP_3024-PP_5558, *oprD*/PP_1207, *erdR, mxtR, infC*, *astA*-II, *topA*, PP_2402-PP_2426, PP_3595-PP_3610, *rpoZ*) that were highly correlated as common mutations that accumulated in response to isoprenol in the TALE experiments (**Figure 2B** and **Supplementary Table 4**). Out of these 12 commonly mutated regions in response to isoprenol, we briefly discuss those mutations that occurred most frequently: *gnuR*, *pcaK*-*opdH*, PP_3024-PP_5558, and *mxtR* (in more than 4 of the 12 evolved isolates, see **Supplementary Table 4**) as major determinants of isoprenol tolerance.

#### Mutations in the Transcriptional Repressor GnuR Suggests its Relevance to Global Stress Responses to Growth Inhibitors

*gnuR* encodes a LacI-family transcriptional regulator (del Castillo et al., 2008) and was commonly mutated. Unique mutations were identified in all isolates from populations collected at all three-time points in the isolates A5-A12, with 100% occurrence from IPL300 and IPL400 backgrounds whereas mutations were identified only in the second and end time points of the isolates from the wild type starting strain A1-A4 (**Figure 2B**, **Supplementary Table 4**). A total of 17 different mutations, including single amino acid changes, small in-/out-of-frame deletions, and early stop codon insertions, were found in its coding sequence (**Supplementary Dataset 1**). This observation suggested that inactivation of GnuR was helpful to achieve high fitness under the isoprenol-induced stress condition. Mutations in *gnuR* were similarly observed in a previous ALE study which tolerized *P. putida* KT2440 against triethylamine hydrogen sulfate, a protic ionic liquid (Lim et al., 2020) having a hydrocarbon chain, but at a less frequency. Its transcript levels were also significantly downregulated (log_2_ fold < −2) when *P. putida* KT2440 was grown in the presence of isopentanol (Lim et al., 2022) implying that the effect of *gnuR* deletion is possibly involved in stress response to various growth inhibitors. Its mutation did not appear to be beneficial to HCHO stress, given no mutations were observed in isolates tolerized against HCHO.

#### Large Deletions in the Evolved Isolates Derived from WT and IPL300 Strains Support Fitness Advantage from Inactivating the Efflux Pump TtgABC

We identified mutations in nearly all the endpoint isolates (11 out of 12 isolates, 92%) either as large deletions ranging from 15-28 kb in the gene locus encoding *pcaK* (PP_1376) to *opdH* (PP_1419) in the evolved lineages from A1-A8 or a small 4 bp deletion and an SNP in PP_1395 or a SNP in PP_1396 (**Figure 2B, Supplementary Figure 2B, Supplementary Dataset 1**) in the evolved lineages from A10-A12. The *pcaK* (PP_1376) - *opdH* (PP_1419) locus contains 45 genes encoding enzymes for the catabolism of aromatic acids (*pcaK*, *pcaF*-I, *pcaTBD*, *pcaC*, *pcaP*, *galP*-II) (Romero-Steiner et al., 1994), efflux systems and transporters (*ttgABC*, PP_1388, *kgtP, tctABC, opdH*), a putative oxaloacetate decarboxylase (PP_1389), a putative acetolactate synthase large subunit (PP_1394), regulators (*ttgR*, PP_1391, PP_1393, PP_1395, *dctD-III*), and several uncharacterized proteins (PP_1390, PP_1391, PP_1392, PP_1393). Given that most of the mutations are gene deletion resulting in loss-of-function (LoF), this implied that inactivating this large genomic locus or specific genes in this locus was beneficial for improving isoprenol tolerance. Notably, mutations in this region were different depending on whether *ttgB* was deleted in starting strain (**Supplementary Figure 2B**). In the evolved isolates derived from the wildtype or IPL300 which harbors an intact *ttgB*, a 10-kb region of *ttgA*-PP_1395 was commonly deleted whereas evolved IPL400 isolates (where *ttgB* had been previously deleted) acquired mutations in only PP_1395 or its neighboring gene (PP_1396). This mutational pattern of losing the function of *ttgB* agrees with our initial observations where IPL400 showed enhanced growth in the presence of 4 g/L isoprenol (**Supplementary Figure 1D**) over IPL300. The effect of PP_1395 is not currently clear, given the lower growth rate improvement by its deletion (see the next section). Nevertheless, taken together, these observations strongly imply that deletion of *ttgABC* correlates with improved fitness in the presence of isoprenol.

#### Large Gene Deletions in the PP_3024-PP_5558 Region Suggest the Expression of These Genes May be Disadvantageous for Cellular Fitness in the Presence of Isoprenol

It was observed that the PP_3024-PP_5558 region was mutated in 6 of the 12 (50%) endpoint evolved isolates (**Figure 2B**). This region includes 53 phage or pyocin-related genes including phage recombinase, LysE-type translocators, an efflux transporter, regulators, a DNA-methylase, proteins coding for contractile tail particles and other hypothetical proteins with unknown functions (Weinel et al., 2002). Five of these strains (lineages A1, A3-A6) identically acquired a large deletion of 40,784 bp whereas the isolate from lineage A12 had a frame-shifting 1-bp deletion only in PP_3024 encoding a hypothetical protein (**Supplementary Dataset 1)**. The commonality in the deletions in the PP_3024-PP_5558 region indicate that expressing these genes could be detrimental to cellular fitness when exposed to isoprenol.

#### Mutations in a Two-Component System Indicate the Suppression of Unnecessary Activation of Acetate Metabolism Genes

MxtR (PP_1695 encoding a sensor kinase MxtR, also called CrbS) was known to consist of a two-component system together with ErdR (PP_1635 encoding a response regulator, also called CrbR) (Henriquez and Jung, 2021). Their coding genes were mutated in five ALE experiments (ALE7 and ALE9-12) and the mutations were mostly observed in endpoint isolates. Only single amino acid change mutations occurred in MxtR (P71S, R603H, L617R), however, a 156-bp deletion or amino acid change mutation occurred in ErdR in ALE11 and ALE12. Previously, this two-component system was reported to activate genes involved in acetate utilization (Henriquez and Jung, 2021); its direct interaction with the promoter regions of *yjcH-actP*-I (PP_1742-3) encoding inner membrane protein and acetate permease, PP_0354 encoding CBS domain-containing protein, *scpC* (PP_0154) encoding propionyl-CoA:succinate CoA transferase were previously observed. Collectively, these observations implicate a potential isoprenol mechanism, suppression of adverse activation of acetate metabolism related genes in the presence of a high concentration of isoprenol.

Overall, DNA resequencing confirmed that 12 gene loci were commonly mutated in response to isoprenol. Several of the evolved isolates had accumulated mutations ranging from SNPs to large deletions in the presence of isoprenol in both the WT and the isoprenol-catabolism deficient backgrounds. These were predominantly observed in regions that encode transcriptional regulators, an efflux pump and non-essential viral proteins (*gnuR*, *pcaK*-*opdH,* PP_1395, PP_3024-PP_5558, *mxtR and erdR*). We propose that these deletions could be major determinants of isoprenol tolerance in *P. putida*.

### 3.4. Validation of a Genetic Linkage to Isoprenol Tolerance and the Impact on Isoprenol Production by Reverse Engineering Common Mutation Targets

We validated the phenotypic linkage of isoprenol tolerance to potentially causal mutations by reverse engineering gene deletions into the WT strain. The four previously discussed regions were chosen for deletion (*gnuR*, *ttgB*-PP_1395, PP_3024-PP_5558, PP_1695) because they frequently accumulated inactivating mutations or large deletions in more than 4 ALE regimes (**Supplementary Table 4**). We also included three specific individual genes (*ttgB*, PP_1395, and PP_3024) for deletion. A total of seven deletion strains were generated. The only significant increase in isoprenol tolerance for this panel of deletion mutants was in the ΔPP_3024 mutant and the larger deletion mutant containing a ΔPP_3024 deletion (i.e., the ΔPP_3024-ΔPP_5558 strain). These two mutants displayed improved growth rates by 1.7-fold on average when grown in the presence of 6 g/L isoprenol M9 media, confirming that the identified mutated regions contributed to the improved tolerance against isoprenol. As the smaller ΔPP_3024 deletion essentially accounts for most of the tolerance improvement (1.6-fold) seen in the larger region mutant. This observation implies that PP_3024 is a major player among the 53 deleted genes in the larger genetic locus. The remaining deletion mutants did show a subtle increase in growth rate but did not meet the threshold for statistical significance with *p* values less than 0.05. Additionally, none of these mutants showed growth on 8 g/L isoprenol (data not shown). Thus, no single mutational event emerged that could enable growth at 8 g/L as seen in the ALE strains (**Figure 2A**).

Further, we investigated the impact of these single gene deletions on isoprenol production by transforming them with the isoprenol biosynthetic plasmid, pIY670 (Banerjee et al., 2024) (**Material & Methods**, **Figure 3B**). This plasmid contains five genes encoding a heterologous isoprenol IPP-bypass pathway. We did not observe any correlation between tolerance enhancement and isoprenol titers. Interestingly, both the large deletions mutants (PP_3024-PP_5558 and *ttgB*-PP_1395) showed a decrease in isoprenol titers compared to the starting strain, with the latter deletion completely abolishing isoprenol production. This finding identified a new set of genes in these regions that are directly (or) indirectly required for isoprenol production. It is also not surprising that the isoprenol titer did not increase above the control strain as the targeted mutations are not implicated in metabolic flux towards isoprenol production and/or transport but are annotated to have ancillary regulatory or unknown functions. Reviewing both the tolerance phenotype and the isoprenol production characterization, the PP_3024 deletion mutant showed significant enhancement in tolerance while retaining heterologous isoprenol production compared to the starting strain and is a candidate for further study. Currently, no information is available for PP_3024, which is annotated as a putative gene with no KEGG orthology or PFam domains (Karp et al., 2019). Although some single mutation events upon reverse engineering could capture the improved tolerance profiles from lab evolution to a certain extent (Lennen et al., 2023; Lim et al., 2020; Mohamed et al., 2020), our data is consistent with isoprenol tolerance behaving as a polygenic trait that relies on the contribution of many different cellular players.

**Figure 3.**
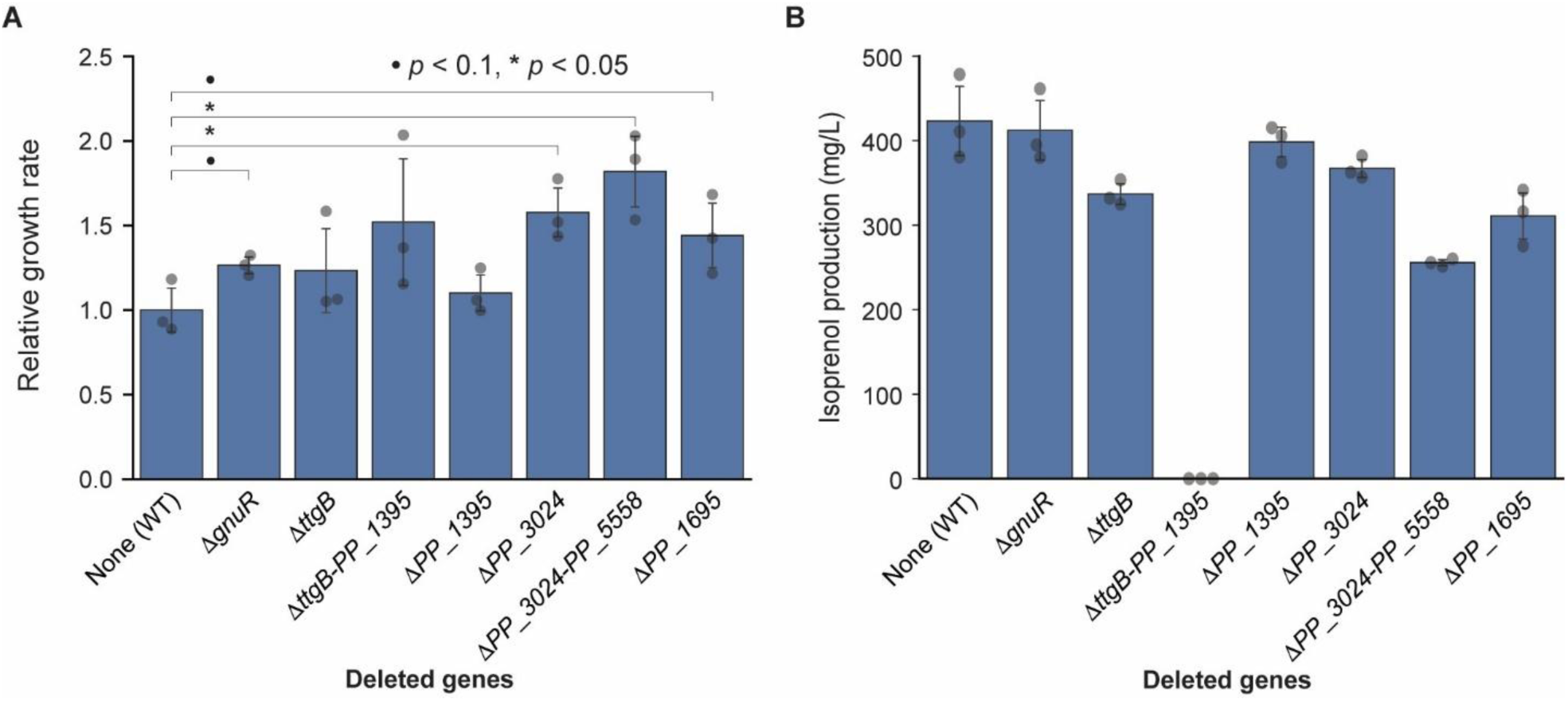
Reverse engineering *P. putida* KT2440 (WT) with commonly mutated targets from the TALE campaign confirmed causality and enabled robust isoprenol production platforms. (A) Relative growth rates of the reverse-engineered strains grown on 4 g/L glucose minimal medium supplemented with 6 g/L isoprenol. (B) Isoprenol production profiles of the reverse engineered strains transformed with the pIY670 plasmid (Banerjee et al., 2024). Transformed strains were grown on glucose (20 g/L) minimal medium with 50 mg/L kanamycin. Cell culture was harvested at 48 h for isoprenol analysis. Values are mean ± SD. • and * indicates *p* < 0.1, *p* < 0.05, respectively.

### 3.5 Global Proteomic Profiling of Evolved Strains Identified Additional Changes Resulting from TALE

We analyzed the global effect of TALE on protein expression in the evolved strains using shotgun proteomics which revealed additional modes complementing the key genetic determinants for isoprenol tolerance. For this analysis, we chose two representative evolved end-point isolates (i.e., A10_F63_I1 and A12_F53_I1) which were derived from the same starting strain (IPL400) but contained mutations in different genes (**Supplementary Table 5**). The two strains showed differential protein expression in about 5% of the total ∼2,400 detected proteins (120 in A10_F63_I1 and 110 proteins in A12_F53_I1) in glucose minimal medium as compared to IPL400.

We then examined the correlation between the observed mutations from TALE and the corresponding protein levels in these two evolved strains. In A10_F63_I1, three (*gnuR*, PP_4063 and *frmA*) out of the total nine mutations, and in A12_F53_I1, four (*gnuR*, PP_4063, PP_3024 and *frmA*) out of the total nine mutations directly translated to concomitant changes in protein levels compared to the starting strain IPL400 (**Figure 4A, Supplementary Table 5**). The remaining mutations were either not detected in the analysis or had no differential protein expression. We observed significant reduction in the protein levels due to frame-shift mutations in PP_3024 compared to the starting strain, reiterating its loss-of-function for enhanced isoprenol tolerance **(Supplementary Note 1 and Supplementary Table 5)** and the consistency between the two - omics analyses. The levels of the transcriptional repressor GnuR (PP_3415) were also reduced in both the evolved strains because of SNPs, confirming its loss-of-function. An intergenic mutation in the region between PP_4061 and PP_4063 also resulted in upregulated levels of the latter coding for a putative long-chain fatty-acid-CoA ligase. This protein has also been implicated in isoprenol catabolism from Rb-TnSeq analysis albeit with a moderate fitness score (Thompson et al., 2020). The other commonly upregulated protein, FrmA (PP_1616) is not directly related to isoprenol tolerance but to the trace levels of formaldehyde in the exogenously added isoprenol during the TALE experiments and is not discussed further. Detailed analysis of the strain-specific proteomics response is described in the supplement **(Supplementary Note 1)**.

**Figure 4.**
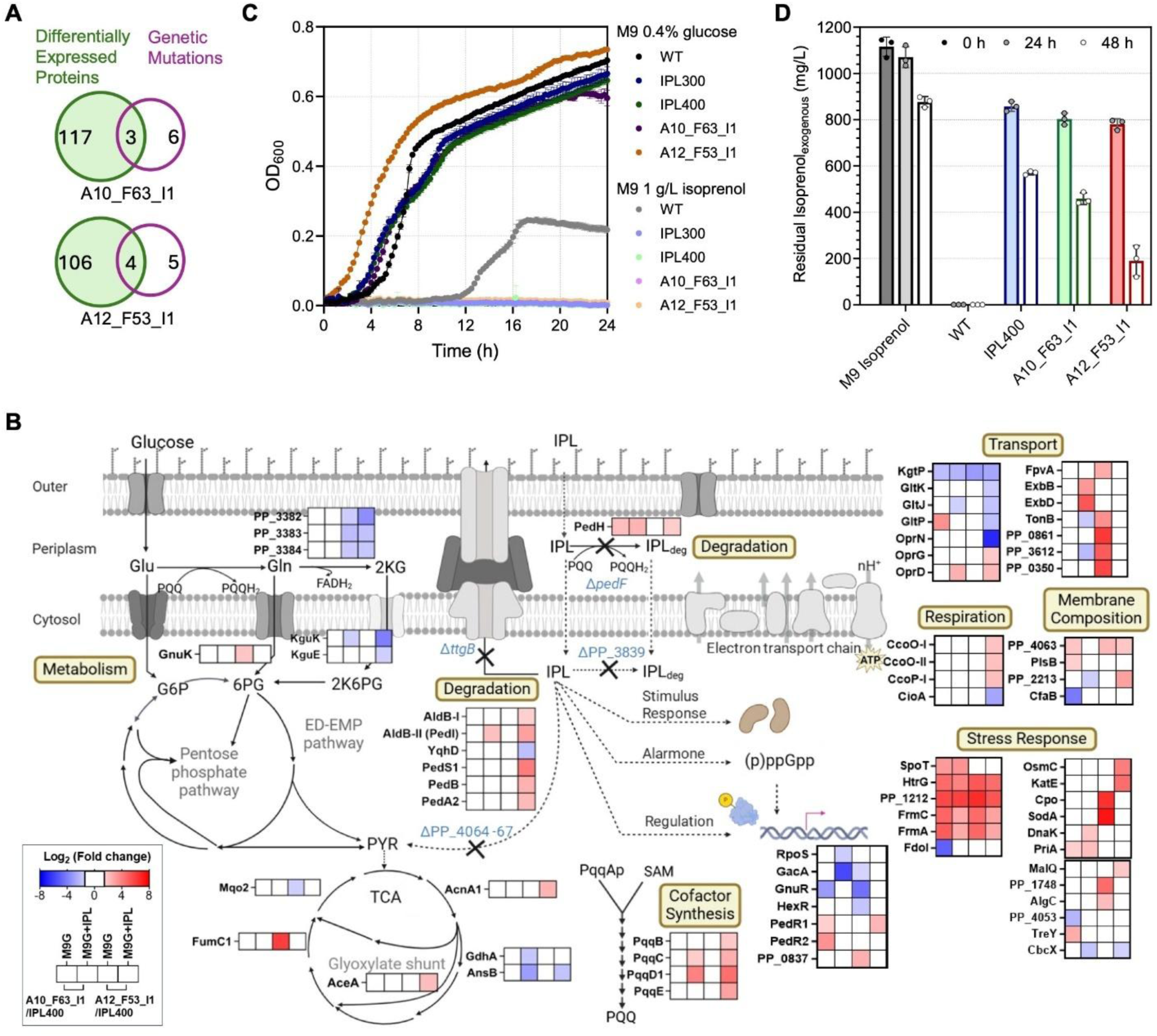
Comparative proteomics of the evolved isolates A10_F53_I1 and A12_F63_I1 and their isoprenol utilization capabilities with respect to the starting strain IPL400. (**A**) Venn diagrams of the number of differentially expressed proteins vs the mutated genes in the evolved isolates. (**B**) Significantly upregulated and downregulated proteins in the metabolic pathways and in various subsystems of the evolved isolates of *P. putida* demonstrating additional mechanisms of isoprenol tolerance. Proteomic profiles of the evolved isolates grown on glucose (4 g/L) minimal medium and supplemented with 4 g/L isoprenol were compared to that of the starting strain IPL400 grown under similar conditions. Significance is defined as log_2_ (Fold change) > ± 1.5, *p* < 0.05. The pathways figure was created in BioRender (Srinivasan, A. (2025) https://BioRender.com/r47f397). **(C)** Growth curves of *P. putida* strains grown on M9 minimal medium with 4 g/L glucose or 1 g/L isoprenol as carbon source (+IP) in 48 microtiter plates for 24 h. Values are mean ± SD of three independent biological replicates. **(D)** Isoprenol degradation capabilities of WT, the IL400 engineered starting strain and the two evolved isolates when grown on co-substrate glucose. Cells grown on M9 4 g/L glucose and ∼1 g/L isoprenol in test tubes were harvested at 24 h and 48 h and the residual isoprenol was measured by GC-FID. Values are mean ± SD of three independent biological replicates.

Interestingly, we also observed differential changes in protein levels coded by genes without detected mutations (**Figure 4A**) similar to the transcriptome response in *E. coli* MG1655 not directly linked to mutations in response to isoprenol stress (Babel and Krömer, 2020). However, in this study, we observed only a few overlapping protein level changes between the two evolved strains and across the growth conditions (with and without isoprenol supplementation) indicating the strains’ divergent responses resulting from TALE although having derived from the same parent (**Supplementary Figure 3A**). These changes include shifts in proteins involved in metabolism, global regulation, transport, cofactor synthesis, activation of stress (or) stringent response and isoprenol degradation.

Significant downregulation of the transcriptional repressors GnuR (in both the evolved strains) and in HexR with concomitant downregulation of the *ptxS* regulon-PP_3382-84 and KguE (in A12_F53_I1). We also observed the significant upregulation of the glyoxylate shunt proteins (AceA and AcnA1) and the high oxygen-affinity cytochrome cbb3-1 type terminal oxidase proteins (such as CcoO-I) in the presence of isoprenol in A12_F53_I1. Both the strains also exhibited upregulation in the co-factor PQQ biosynthesis proteins (PqqB-E). Collectively, these imply enhanced glucose metabolism via the ED-EMP pathway and the cellular response to high energy demands for solvent tolerance in the evolved strains.

The evolved strain A10_F63_I1 showed upregulation of a primosome assembly protein (PriA), chaperone protein (DnaK) and exhibited a stringent response by upregulating SpoT/PP_5302 involved in (p)ppGpp biosynthesis on glucose minimal medium. Upon isoprenol supplementation, we observed downregulation of RpoS and GacA thereby indicating a global regulatory effect in play in this strain. On the other hand, A12_F53_I1 showed significant upregulation in proteins involved in the oxidative stress response such as a hydroperoxidase (KatE), Chloroperoxidase (Cpo), Superoxide dismutase (SodA), osmotic stress response (OsmC, PP_4707), biosynthesis of osmoprotectants such as trehalose, glycogen and NAGGN (MalQ, PP_1748, AlgC) and upregulation of the flagella biosynthesis regulator YcfJ (PP_0837) on glucose minimal medium or in the presence of isoprenol. We also observed changes in the levels of the proteins involved in composition, activation or modification of fatty acids and phospholipids (PP_4063, PlsB, PP_2213, CfaB) in addition to transporters including outer membrane porins (OprG, OprD) and those involved in iron/siderophore uptake (such as FpvA, ExbBD/TonB) in both the strains. Two of the proteins that were constitutively highly upregulated in both the strains include FmdB (PP_1212) and the signal transduction protein HtrG (PP_3631). The former protein has been observed to be important for tolerance to 1-butanol in *P. putida* BIRD-1 (Cuenca et al., 2016). Notably, several proteins involved in alcohol degradation (such as PedH, AldB-I, AldB-II, PedS1, PedA2) were upregulated in both the evolved isolates mostly in the presence of isoprenol. This observation suggested the possibility of alternate routes to isoprenol degradation which could have been activated by evolution and were now contributing to enhanced stress tolerance in the presence of a co-substrate regardless of their inability to utilize isoprenol as the sole carbon source (**Figure 4C**). Consistent with most upregulated proteins in this subsystem, the evolved strain A12_F53_I1 showed relatively faster degradation of exogenously added isoprenol compared to the IPL400 starting strain (**Figure 4D**). Nevertheless, the isoprenol degradation capabilities of the other evolved strains (derived from IPL300 and IPL400) were significantly lower compared to the wildtype strain or its evolved derivatives wherein active catabolism also potentially enhances tolerance (**Supplementary Figure 3B**).

Thus, TALE experiments with the stressor isoprenol cause mutations with both direct and indirect effects which could be elucidated by functional genomics approaches and physiological studies. These changes are consistent with well-documented solvent response of *P. putida* involving upregulation of DNA-repair systems, chaperones to refold denatured proteins, activation of oxidative and osmotic stress response including changes in membrane fluidity caused by adjustments in fatty acid and lipid composition and in some cases, active degradation of the target compound (Bojanovič et al., 2017; Ramos et al., 2015).

### 3.6 Rational Engineering of Evolved Strains to Restore Glucose Consumption and Isoprenol yields

To examine the applicability of the evolved strains for use as isoprenol production chassis, we introduced the isoprenol production plasmid, pIY670 and examined their isoprenol production. Initially, all the evolved strains showed an unexpected overall reduction in isoprenol titers regardless of the starting strain background (**Figure 5A**) which was contrary to the generally maintained titers in the reverse engineered WT strain with the single gene deletions (**Figure 3B**). These decreases were likely linked to significantly reduced consumption (by half) of glucose (**Figure 5B**). Although this suggested an overall trade-off between isoprenol production and tolerance upon TALE, the variability in degree to which isoprenol titers were reduced did not strongly correlate with the tolerance levels of these different isolates. This observation may be attributed to the polygenic nature of isoprenol tolerance, probable alterations in membrane properties, or a heightened metabolic burden following the transformation of the evolved strains with the isoprenol pathway plasmid. Among the evolved production strains, those engineered from A9_F65_I1 and A10_F63_I1 produced relatively higher concentrations of isoprenol (60–70 mg/L) at 48 hours compared to the other strains. However, this production remains significantly lower than their starting strain, which produced 280–400 mg/L at the same time point (**Figure 5A**).

**Figure 5.**
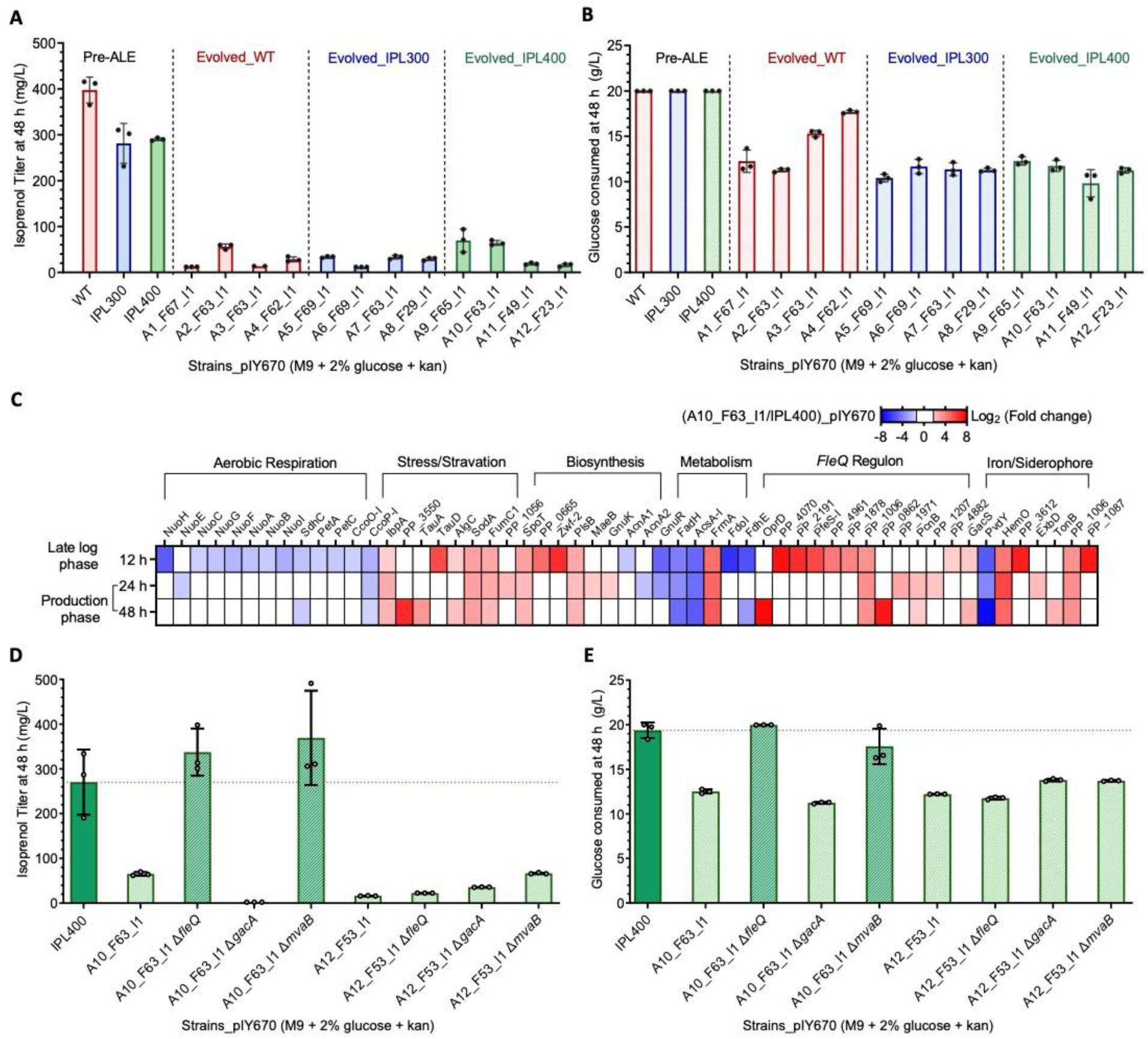
Rational engineering improved isoprenol titers in the evolved strain A10_F53_I1. **(A)** Isoprenol titers (mg/L) in the wild type, starting and evolved isolates of *P. putida.* The strains transformed with the isoprenol production plasmid pIY670 (see Materials and Methods). Cells were harvested at 48 h for analysis. Values are mean ± SD. **(B)** Residual glucose (g/L) measured after 48 h in the cell supernatants. **(C)** Proteomic analysis of the evolved strain A10_F63_I1 expressing pIY670 compared against the starting strain IPL400 expressing pIY670, harvested at 12 h (Late growth phase), 24 h and 48 h (Production phase). Significantly upregulated and downregulated proteins in various subsystems were plotted as a heatmap. Significance is defined as log_2_ (Fold change) > ±1.5, p<0.05. **(D)** Isoprenol titers (mg/L) in the rationally engineered strains, evolved strains A10_F63_I1 and A12_F54_I1 vs. the starting strain IPL400. Markerless deletion of selected gene targets (*fleQ, mvaB,* and *gacA,* see Materials and Methods) identified using proteomics and computational aided metabolic engineering (Banerjee et al., 2024) was carried out. **(E)** Residual glucose (g/L) measured in the cell supernatants from the production runs harvested at 48 h. Values are mean ± SD.

Shotgun proteomics using a representative pre- and post-TALE strain harboring the isoprenol production plasmid helped us to further understand the limitations of heterologous isoprenol production (**Figure 5C**). We sampled the production strains, A10_F63_I1_pIY670 and IPL400_pIY670 at late log phase (12 h) and in the production phase (24 h and 48 h). Firstly, the low titers in this evolved production strain were not due to enhanced isoprenol degradation as we did not observe differential expression of alcohol dehydrogenases except for overexpression of CalB at 24 h. Secondly, it was not due to the poor expression of the heterologous production pathway in A10_F63_I1_pIY670 (**Supplementary Figure 3C**). We then observed a significant reduction in the aerobic respiration pathway proteins (e.g., NuoABCEFGHI) and an increase in stress-related proteins (e.g., SpoT, SodA, IbpA) in the evolved production strain. These changes in the global proteome indicated a stringent condition, which is consistent with the impaired glucose consumption. Notably, the highest number of significantly changed proteins (**Supplementary Table 6**, 35 genes of the 478 proteins, e.g., OpoD, PfeS-I, GacA) were found to be regulated by the global regulator *fleQ*, involved in flagellar and motility, adhesion, and exopolysaccharide production. The abundance of many of these proteins increased, potentially suggesting their undesired activation (**Figure 5C**).

We therefore reasoned that the deletion of *fleQ* would help us in leveraging the metabolic plasticity inherent to these strains overcoming any unanticipated unfavorable cellular rewiring to remedy the loss in isoprenol titers in the evolved isolate A10_F63_I1_pIY670. Previously, the deletion of *fleQ* has been shown to aid heterologous production by reallocation of cellular resources (Blanco-Romero et al., 2018; Kim et al., 2024), supporting our hypothesis. Indeed, the evolved isolate A10_F63_I1 Δ*fleQ_*pIY670 showed enhanced glucose uptake (to exhaustion at 48 h) compared to a residual glucose value of 7.5 g/L at the same time point in the starting strain thus enhancing the overall isoprenol yields (**Figure 5E**). The isoprenol titers also significantly increased by 4.8-fold (337 mg/L) at 48 h (**Figure 5D**).

Additionally, rational engineering *via* a previous genome scale metabolic model guided flux analysis also helped restore isoprenol titers in the evolved engineered strain (Banerjee et al., 2024). To this end, we deleted *mvaB* to prevent the reverse reaction (blocking isoprenol accumulation) from 3-hydroxy 3-methylglutaryl CoA to acetoacetyl CoA. Although we did not see any differential expression of MvaB in the production strain (**Supplementary Figure 3**), we observed that the A10_F63_I1 Δ*mvaB_*pIY670 strain now produced 370 mg/L of isoprenol up from 70 mg/L (**Figure 5D**). However, this did not restore complete glucose utilization at 48 h (**Figure 5E**) suggesting that deletion of *mvaB* alone is not sufficient to rewire the impeded carbon metabolism back to pre-ALE levels.

Thus, rational engineering of the tolerized isolate A10_F63_I1 resulted in engineered strains that were now tolerant to up to 8 g/L isoprenol whilst producing significant isoprenol titers as the starting strains. On the other hand, none of the deletion strains from the A12 lineage showed any dramatic improvement in isoprenol titers (**Figure 5D**). This further substantiated our observations from the proteomics datasets in the prior section that these two isolates exhibited differentially in response to an exogenous isoprenol stress.

## 4. Conclusions

Enhancing microbial tolerance is a necessary approach when developing efficient biofuel production processes. When isoprenol is converted to the bio-jetfuel DMCO, our recent technoeconomic analysis for commercial viability of this process demands that isoprenol be generated from lignocellulosic sugars at 90% theoretical yield or higher (Baral et al., 2023, 2021). Using current growth conditions and yield calculations (Banerjee et al., 2024), this would be equivalent to producing equal or greater than 6.2 g/L isoprenol from 20 g/L glucose. In this study, we successfully applied the TALE approach to enhance the tolerance of *P. putida* KT2440 and its genetically engineered derivatives to isoprenol, achieving stable growth at concentrations as high as 8 g/L. At first glance, this would satisfy the criteria to be useful for microbial isoprenol bioconversion and even surpasses the 6.8 g/L limit reported in a study in another gram-negative host, *E. coli* K-12 MG1655 (Babel and Krömer, 2020). Integrating process strategies such as two-phase extraction systems or *in-situ* alcohol removal could further improve viability and scalability to meet productivity targets from complex carbon streams. Thus, the TALE approach was successful in meeting a tolerance benchmark related to economic feasibility.

This work provides a comprehensive experimental mutational landscape to identify key genetic determinants of isoprenol tolerance by physiological and genomic analyses from three starting genotypes. One finding was increased isoprenol degradation in a subset of evolved lineages. However, growth analysis on isoprenol and/or with a co-substrate glucose in the starting strains and some of the evolved isolates showed that the catabolism of isoprenol as a carbon source in *P.putida* was not always particularly relevant to the resistance phenotype, unlike ethanol in *E.coli* (Goodarzi et al., 2010). In addition, only one of the evolved isolates (A12_F53_I1) showed significant enhancement (67%) in degrading isoprenol to a yet-unknown product over its catabolism-mitigated starting strain IPL400. Other characterized isolates had less of an increase in degradation. While no mutations in catabolic enzymes were detected from whole genome resequencing to support this claim, comparative proteomics analysis pinpointed several upregulated enzymes (such as AldB-I, AldB-II and PP_2668) that may have promiscuous alcohol degradative activity towards isoprenol in this evolved strain.

Mutations in *gnuR*, *ttgB*-PP_1395 and PP_3024-PP_5558 in the evolved strains resulted in clear loss-of-function mutations and these findings were supported by a concomitant decrease in protein levels. Some intergenic mutations such as in PP_4063 could have resulted in enhancing mutations (supported by increased protein abundance) while the effect of others (such as SNP in PP_1695) could not be verified based on our current analysis and warrants further characterization based on activity assays. One of the most notable outcomes of our study is the improved isoprenol tolerance via inactivation of TtgB in strains where it was originally functional. Contrary to many efflux pump studies such as that of AcrB in *E.coli (Fisher et al., 2014)* which demonstrated improved susceptibility (and thereby reduced tolerance) to solvents upon its deletion, our observation highlights the genetic plasticity of gram-negative microbes to alcohol tolerance, as the efflux pump TtgB has also been implicated in enhanced tolerance to isoprenol and n-butanol in *P. putida* (Basler et al., 2018; Cuenca et al., 2016). Although reverse engineering (via generation of isogenic deletions in the WT strain) indicated a combinatorial effect of multiple mutations to achieve the tolerance limits observed through TALE, deleting PP_3024 and *ttgB* in the chassis strains demonstrating gram-scale production (Banerjee et al., 2024) could be a rational next step. In addition, the isoprenol tolerance mechanisms identified here using genomics showed little overlap with those observed in the *E. coli* study (Babel and Krömer, 2020). While in the latter strain, isoprenol tolerance was linked to mutations in genes associated with membrane composition, integrity and activation of stress response, mutations in *P. putida* inactivated efflux pumps and undesired genes.

While the increased isoprenol tolerance in evolved strains generated in this study was promising, evolved strains did not show a corresponding benefit towards producing isoprenol when transformed with a heterologous pathway. However, guided by proteomics analysis and prior research, we restored isoprenol production levels by deleting the biofilm and flagella master regulator, *fleQ* and *mvaB* respectively, yielding a strain that was both tolerant to isoprenol and similarly functional as a chassis compared to the starting strain for isoprenol production. Ideally, the mutations enriched by TALE would engender non-degradative mechanisms for product tolerance. There are some exemplary ALE-driven reports where this duality is observed - for native yeast ethanol production (Mavrommati et al., 2023), L-serine production in *E. coli* (Mundhada et al., 2017) or for heterologous isobutyrate and 2,3-butanediol production in *E. coli* (Lennen et al., 2023). However, given the indication that isoprenol degradation is a possible tolerance mechanism, further iterative engineering cycles are likely warranted to understand the genetic basis of the enhanced degradation to eliminate it, as this tolerance mechanism is antagonistic to production.

In summary, the TALE approach successfully resulted in robust isoprenol tolerized strains. The enhanced tolerance can be significantly advantageous to further develop chassis with superior titers, rates, and yields, which are key factors in determining overall commercial viability. Ultimately, the development of such resilient strains can accelerate the transition to sustainable aviation fuels by making biobased isoprenol production more economically feasible. As such, this advancement represents a key step toward complementing the use of fossil fuels and mitigating the environmental impact of the aviation sector.

## Acknowledgments

This work was part of the DOE Joint BioEnergy Institute (http://www.jbei.org) supported by the U.S. Department of Energy, Office of Science, Office of Biological and Environmental Research, through contract DE-AC02-05CH11231 between Lawrence Berkeley National Laboratory and the U.S. Department of Energy. The work was also supported by the U.S. Department of Energy Bioenergy Technologies Office. The United States Government retains and the publisher, by accepting the article for publication, acknowledges that the United States Government retains a nonexclusive, paid-up, irrevocable, worldwide license to publish or reproduce the published form of this manuscript, or allow others to do so, for United States Government purposes. H.G.L appreciates the National Research Foundation of Korea (NRF) grants funded by the Korea government (MSIT) (grant number RS-2024-00334792 and RS-2024-00399277). We also thank Marc K. Abrams at University of California San Diego for manuscript editing.

## Author Contributions

Conceptualization: HGL AS TE AM AF; Data curation: HGL AS; Formal analysis: HGL AS TE AM AF; Funding acquisition: AM AF; Investigation: HGL AS RM ISY MHN MW YC JWG; Methodology: HGL AS RM ISY TE; Project administration: TE AM AF; Resources: BOP TSL CJP TE AM AF; Software: HGL CJP; Supervision: TE AM AF; Validation: HGL AS TE; Visualization HGL AS; Roles/Writing - original draft: HGL AS TE; and Writing - review & editing HGL AS TE AM AF. All authors read, gave feedback, and approved the submission of the manuscript for publication.

## Supplementary Material

### Supplementary Note

#### Supplementary Note 1. Detailed Proteomics Characterization of the Evolved strains

##### Protein Levels That Directly Correlated to Mutations Acquired during TALE in the Evolved strains A10_F63_I1 and A12_F53_I1

In both the evolved strains, some mutated genes were differentially expressed under standard laboratory growth conditions (without exogenous isoprenol addition, **Supplementary Table 5**). Not all mutants could be characterized due to inherent limitations in the shotgun proteomics detection method. Several proteins of note are described here. The basic amino acid porin encoded by *oprD* (PP_1206) with a C→T SNP in the intergenic oprD/PP_1207 (-360/-331) region showed a significant increase in protein levels in the presence of isoprenol in both the evolved strains. As briefly mentioned in the resequencing section earlier, the intergenic region between *frmA* and PP_1615 also contained a SNP; correspondingly, we observed a constitutive increase of FrmA protein expression in both strains. The transcriptional repressor GnuR (PP_3415) showed significantly lower protein levels on glucose minimal medium. This downregulation could be related to changes in protein activity from two substitution mutants (V46I and G167R). The V46I mutation (in isolate A12_F53_I1) lies in a probable DNA binding domain and G167R (in isolate A10_F63_I1) lies in a substrate recognition domain (Finn et al., 2011). We also observed significant constitutive downregulation in the AraC family transcriptional regulator PP_1395 in isolate A12_F53_I1, where genome resequencing indicated point mutation corresponding to an L81R amino acid mutation in a putative ligand binding domain could change protein stability or abundance. Finally, in the same isolate, a 1-bp deletion in the coding region at basepair 167 in the PP_3024 coding sequence would result in an early ribosome termination where the frameshift mutation would generate a premature stop codon. These analyses corroborated the downstream impact on protein levels as hinted by the DNA resequencing, but the change in protein abundance could not be predicted by the sequence information alone.

##### Additional proteomic responses in the evolved strains in response to isoprenol stress

Upon isoprenol supplementation, ∼8% of the detected proteins were differentially expressed in both the evolved strains compared to IPL400 (**Supplementary Figure 3A**). This clearly indicated proteomic shifts in response to isoprenol stress which could be either direct or indirect consequences of mutations resulting from TALE. The majority of differentially abundant proteins were unique to each isolate. Specifically, the isolate A10_F63_I1 has a significant upregulation of the stringent response regulator SpoT/PP_5302 (on both minimal medium and upon isoprenol supplementation). In this isolate, the master regulator of stringent response involved in (p)ppGpp synthesis, RelA was not affected in its gene sequence (**Figure 2B**) or in its protein levels but the upregulation of its complement SpoT (which is a (p)ppGpp hydrolase with a weak synthase activity) was observed. This could induce large-scale transcriptional changes leading to activation of various stress response regulons. In addition, (p)ppGpp directly inhibits other enzymes thereby reallocating cellular resources towards adaptation and survival under stressful conditions (Gaca et al., 2015). On the other hand, the two-component system regulator GacA and its regulon, the sigma factor RpoS, were both significantly downregulated only in the presence of isoprenol in this isolate.

While the isolate A12_F53_I1 showed more downregulated genes than A10_F63_I1, they could not be easily categorized into subsystems. We thus focused on the upregulated genes in this isolate. Many alcohol metabolism related proteins such as dehydrogenases, two component signaling systems, associated transporters and associated cofactor synthesis proteins were differentially upregulated in the presence of isoprenol. Apart from PedH, AldB-II and OprD that were also observed in A10_F64_I1, we also observed upregulation of the aldehyde dehydrogenase AldB-I (**Figure 4B**). It was also interesting to note the significant upregulation of the isoprenol responsive two-component system sensor histidine kinase/response regulator PedS1 /PP_2664 in this isolate. In addition, PedB and PedA2 are ABC transporters reported to be involved in efflux of chloramphenicol, aromatics and more recently lanthanides (Fernández et al., 2012); (Wehrmann et al., 2019) and were upregulated in this isolate. Transposon mutants in coding sequences for these proteins also have negative fitness when grown on isoprenol (Thompson et al., 2020) suggesting that they could facilitate efflux or transport of isoprenol (complementing for the loss of *ttgB*) or contribute to stabilize membrane architecture with other upregulated outer membrane proteins (OprD, OprG).

## Supplementary Figures

**Figure S1.**
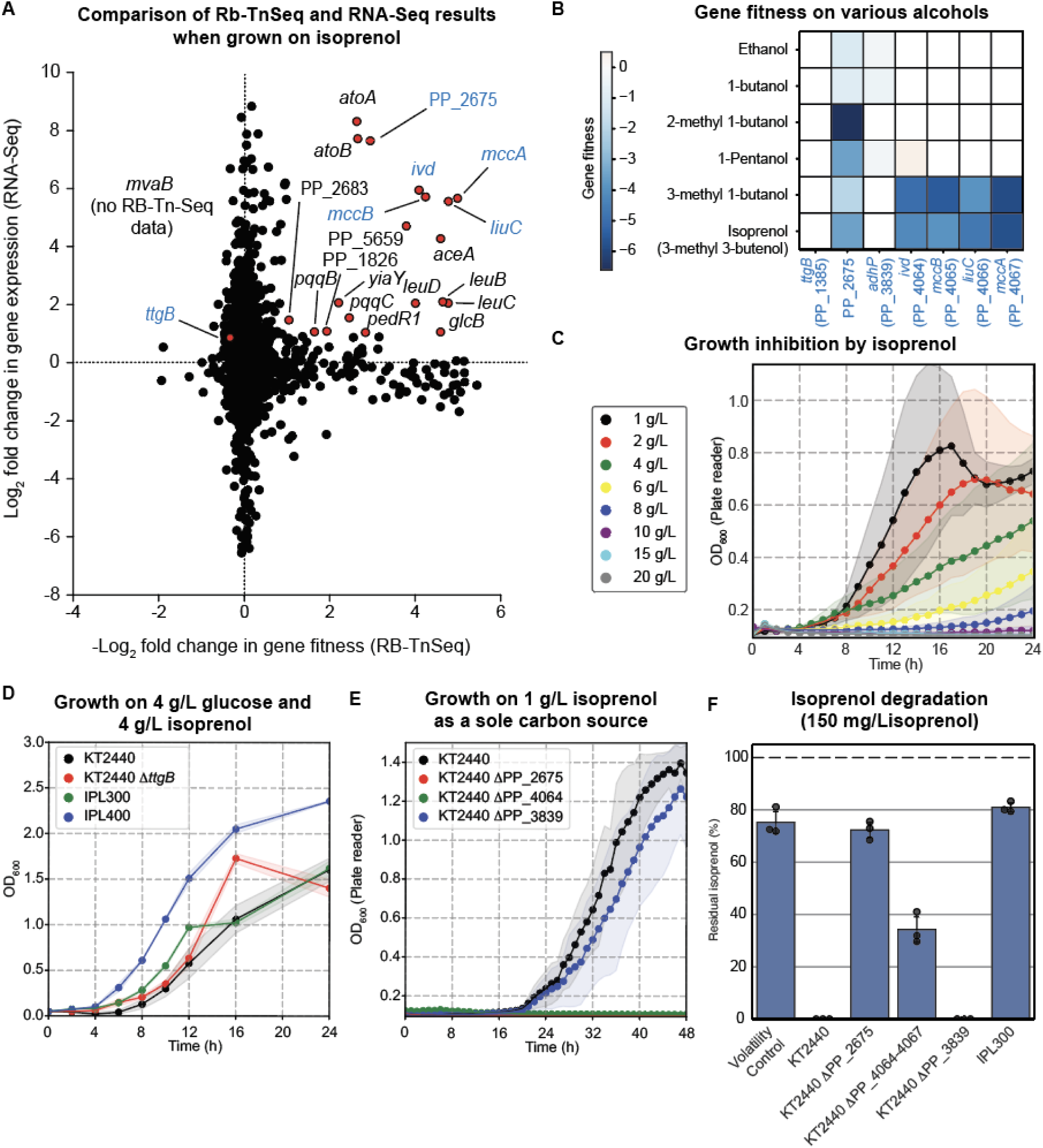
Construction of the starting strains for tolerization ALE experiments. (**A**) A scatter plot of gene fitness from RB-TnSeq and gene expression from RNA-Seq when WT cells were grown on isoprenol as a sole carbon source in a minimal M9 condition. Positive values on the *x*-axis indicate a fitness defect, and the *y*-axis indicates a fold change of gene expression. A gene fitness for each gene is obtained by dividing the abundance of a gene in cells grown on isoprenol by the abundance in cells grown on glucose. A gene expression fold change was similarly calculated by using DESeq2 (Love et al., 2014) using the glucose condition as a reference. (**B**) Rb-TnSeq fitness profiles (on various alcohols) of the genes targeted for deletion to build the strains IPL300 and IPL400 for TALE. (**A** and **B**) Common genes were colored in blue. (**C**) Growth of *P. putida* on M9 minimal medium supplemented with 4 g/L glucose and varying concentrations of isoprenol (0-12 g/L) in a microtiter plate. (**D**) Growth of WT and engineered strains IPL300 and IPL400 in M9 minimal medium with 4 g/L glucose supplemented with 4 g/L isoprenol grown in flasks. (**E**) Growth of the wild-type and engineered *P. putida* KT2440 strains in the M9 medium supplemented in 1 g/L isoprenol as a sole carbon source. (**F**) Percentage of remaining isoprenol after incubating cells for 48 h in the presence of 150 mg/L isoprenol in M9 minimal medium.

**Figure S2.**
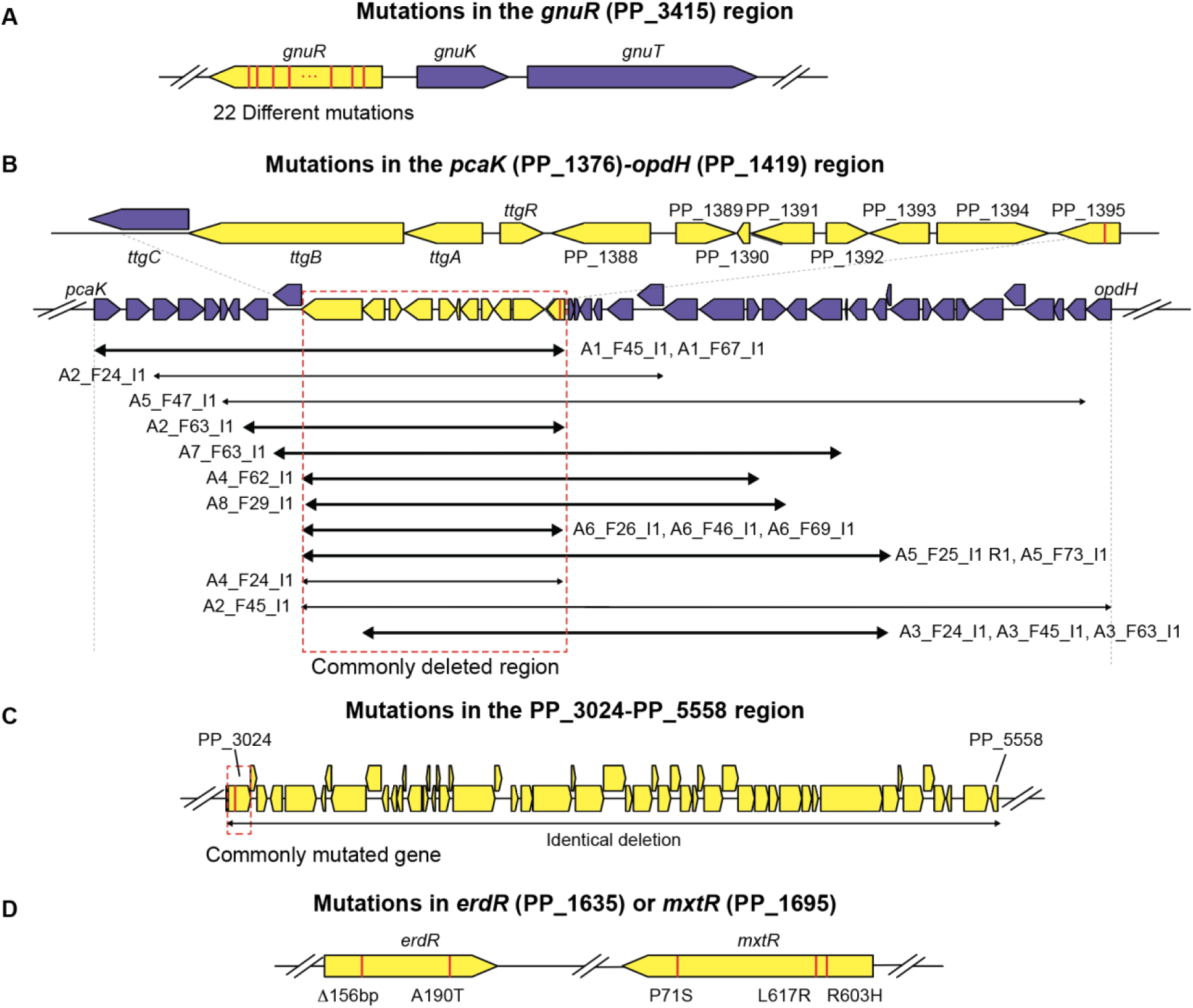
Common mutations from isoprenol TALE identified in the evolved strains. Mutated regions are colored yellow. Mutations are highlighted in red as vertical bars.

**Figure S3.**
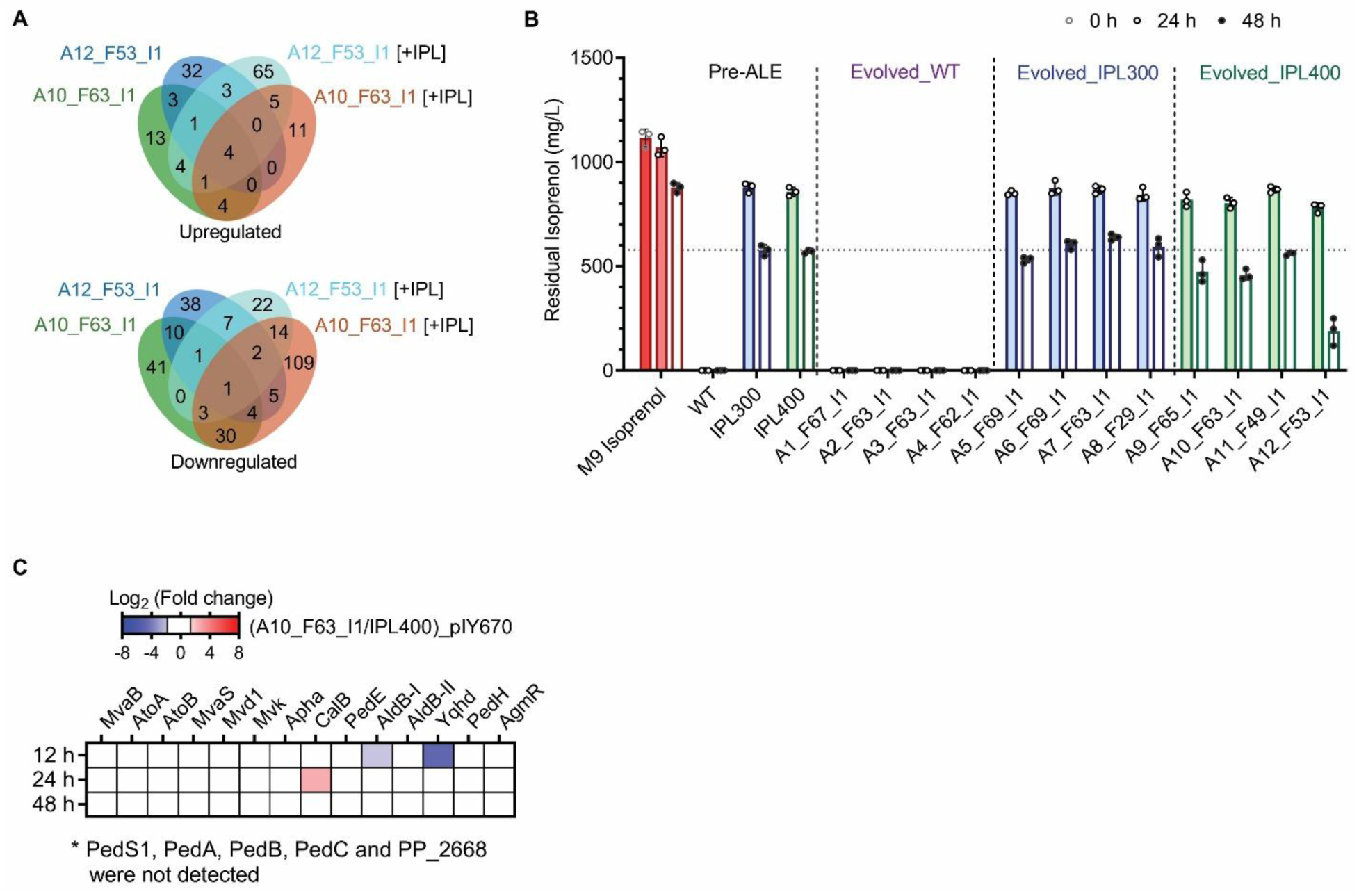
Proteomics analysis of selected evolved strains and their isoprenol degradation capabilities. (**A**) Venn diagram shows the number of common and uniquely up and downregulated proteins in the evolved strains on both glucose minimal media and that supplemented with 4 g/L isoprenol (+IPL). (**B**) Isoprenol degradation capabilities of WT, the starting strains IPL300and IPL400 and the evolved strains when grown on co-substrate glucose. Cells grown on M9 4 g/L glucose and ∼1 g/L isoprenol in test tubes were harvested at 24 h and 48 h and the residual isoprenol was measured by GC-FID. Values are mean ± SD of three independent biological replicates. (**C**) Differential protein expression of the isoprenol pathway proteins and candidate isoprenol degradation enzymes in the evolved strain A10_F63_I1 expressing pIY670 compared against the parent strain IPL400 expressing pIY670, harvested at 12 h, 24 h and 48 h. Significantly upregulated and downregulated proteins in various subsystems were plotted as a heatmap, Significance is defined as log_2_ (Fold change) > ±1.5, p < 0.05.

**Supplementary Table 1.**
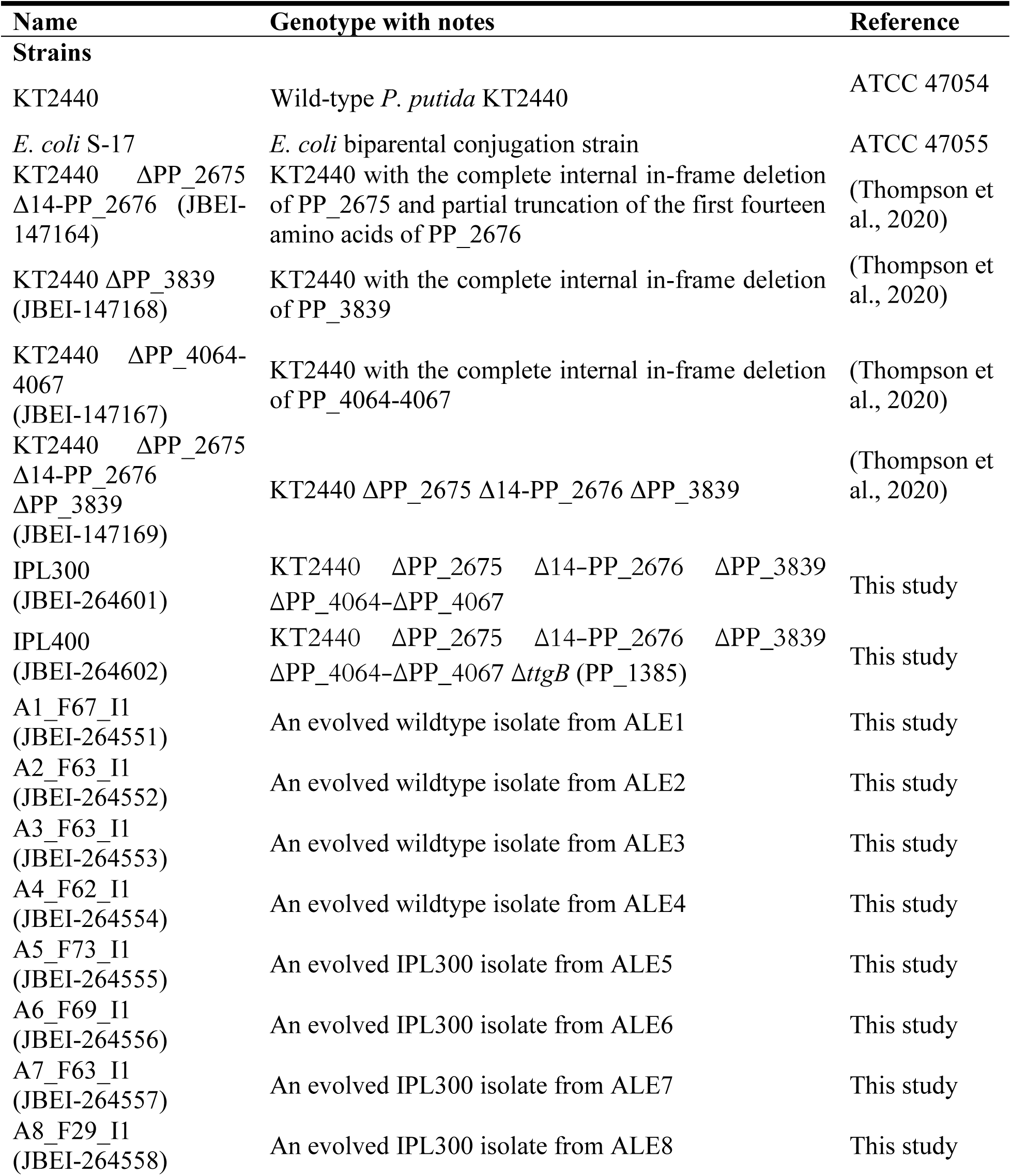
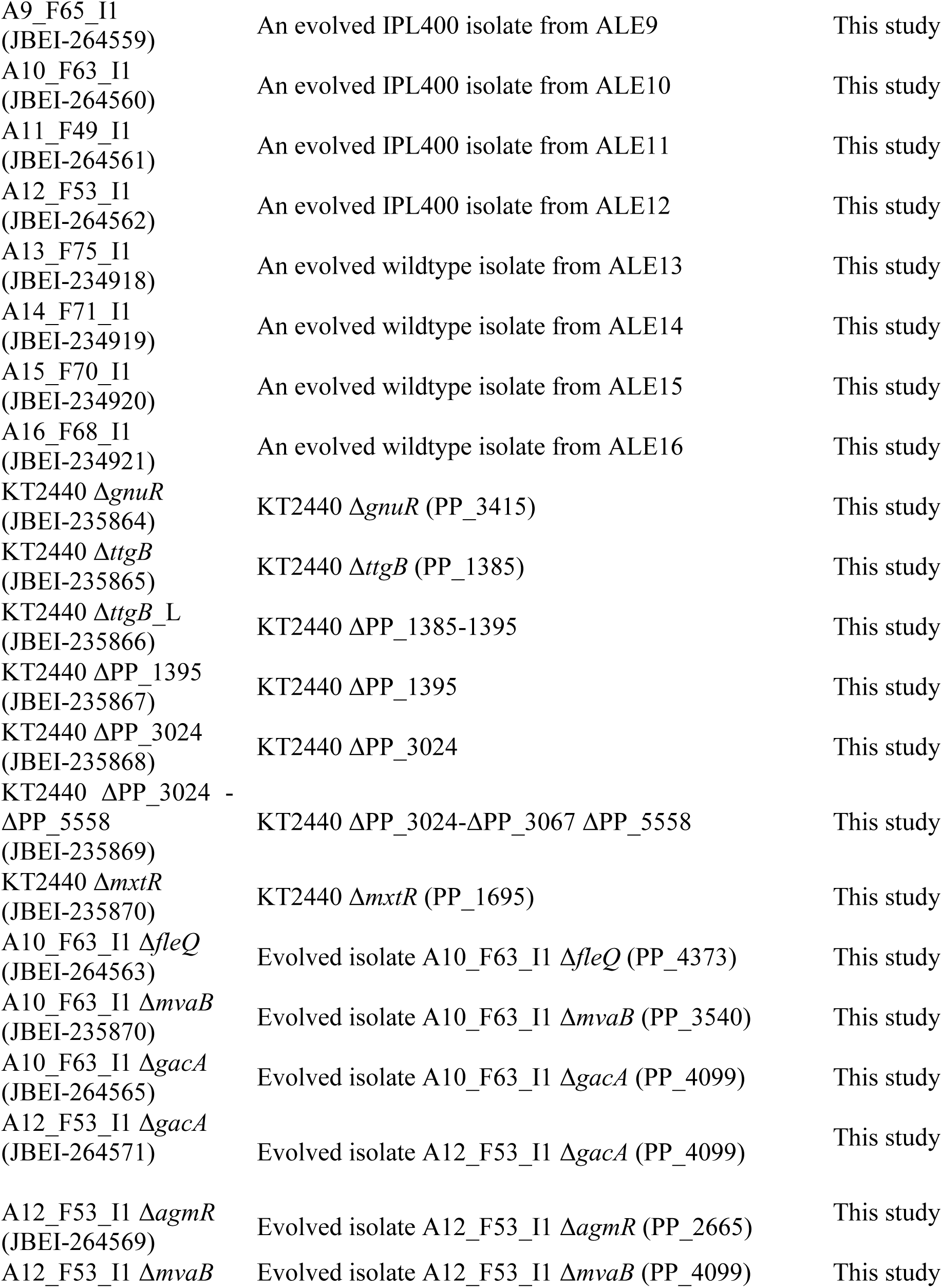
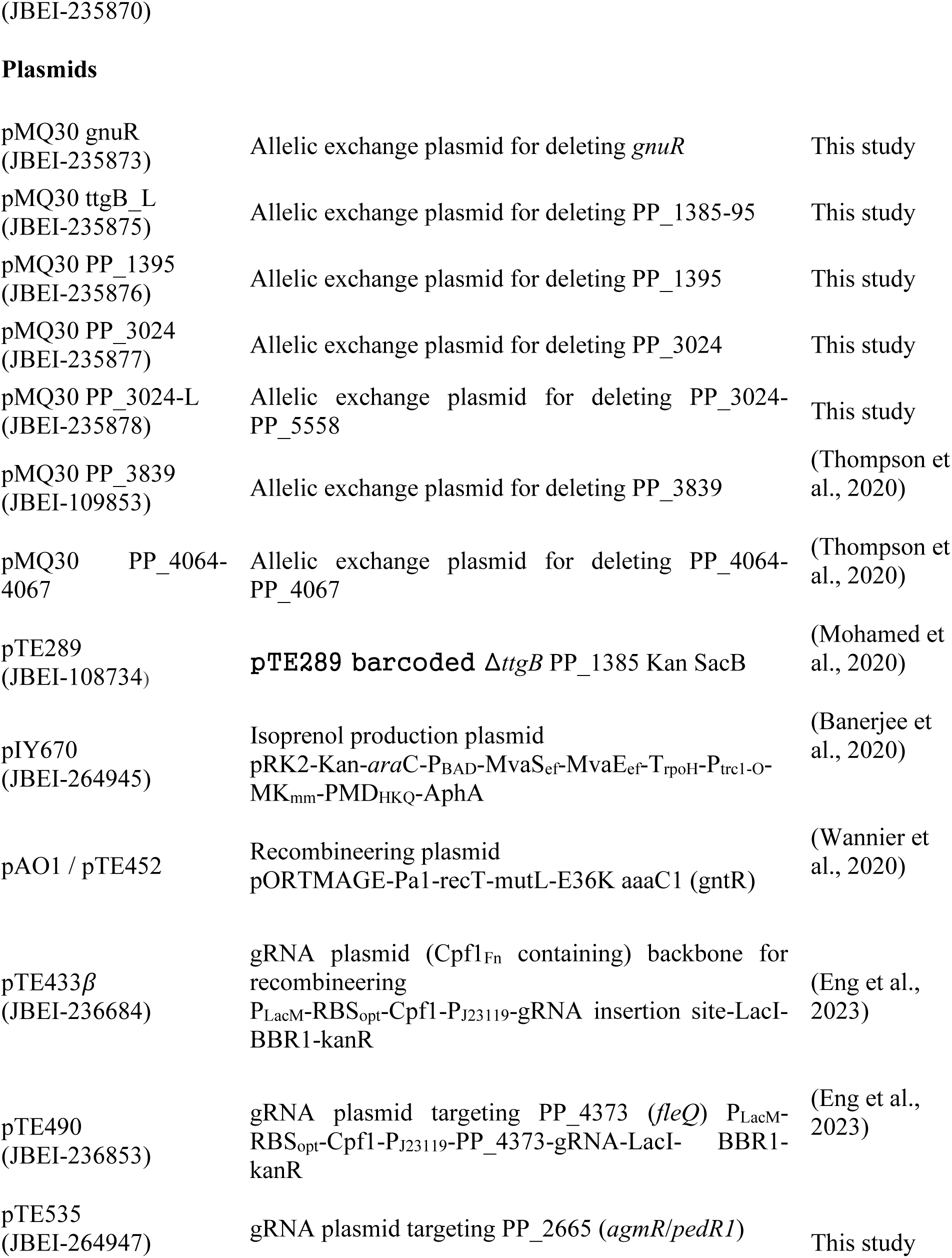
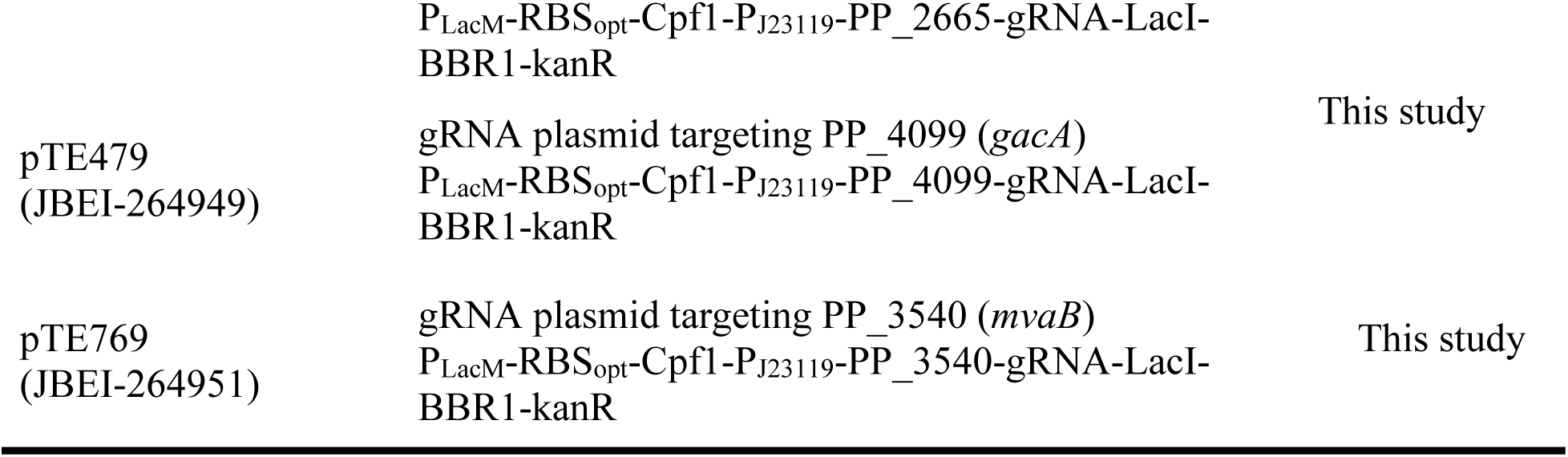
Strains and plasmids used in this study.

**Supplementary Table 2.**
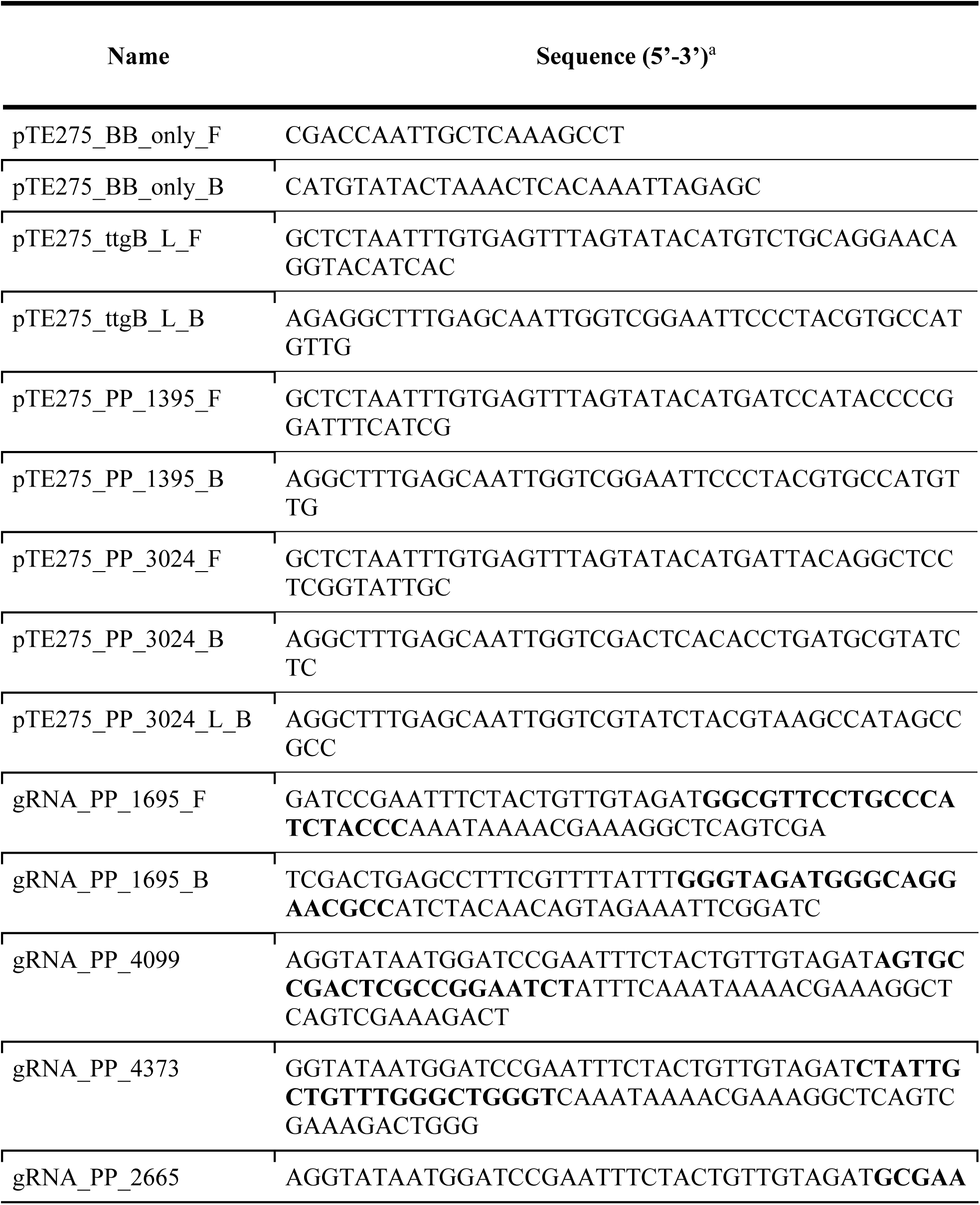
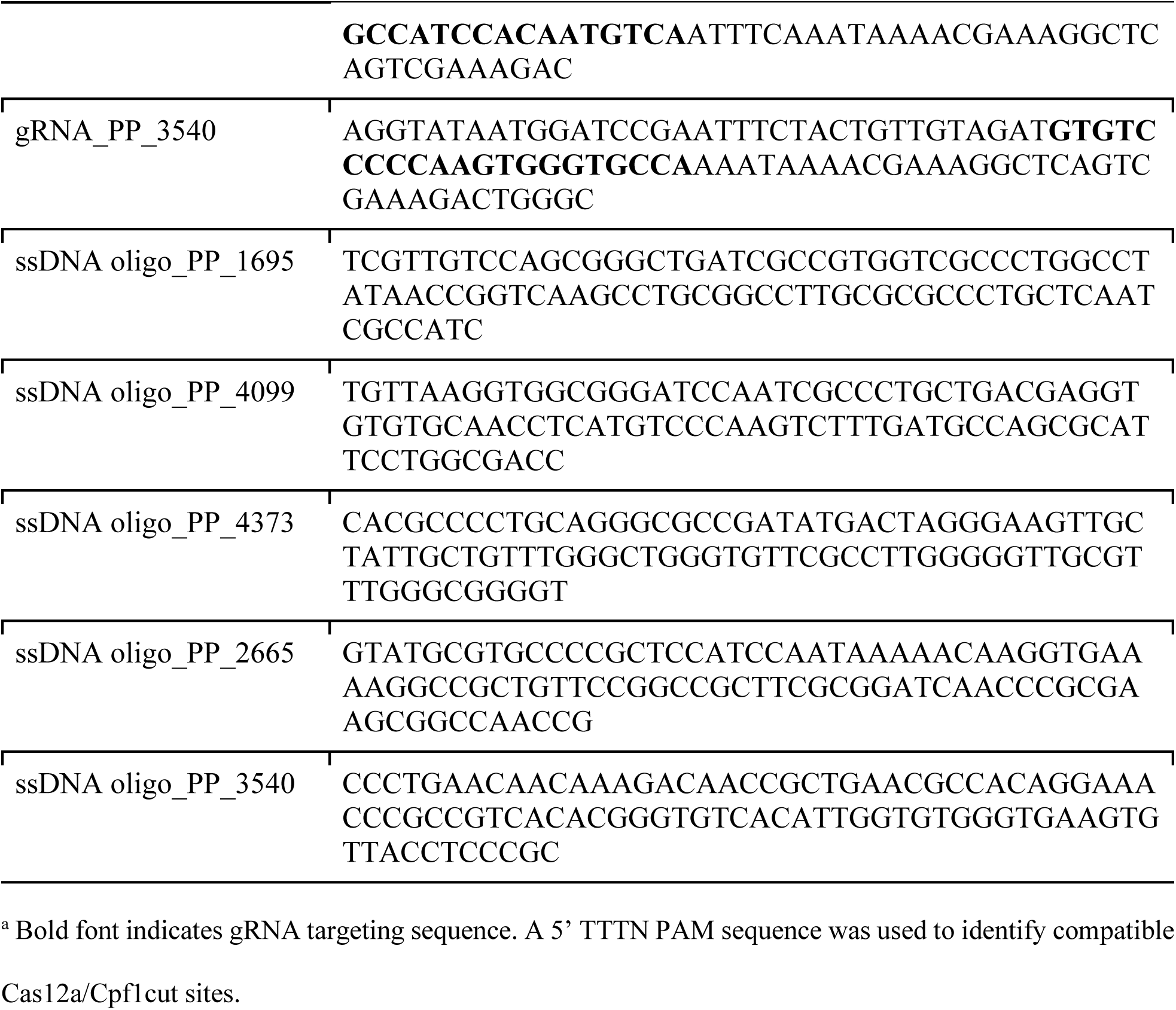
Oligonucleotides used in this study.

**Supplementary Table 3.** Summary of the Tolerization ALE (TALE) to isoprenol.

|  | ALE# | Starting concentration | Initial growth rate (h<sup>-1</sup>) | Ending concentration | Final growth rate<sup>b</sup> (h<sup>-1</sup>) | Passages # | Generations | CCD (10<sup>12</sup>) |
| --- | --- | --- | --- | --- | --- | --- | --- | --- |
| **Isoprenol TALE KT2440** | 1 | 4 g/L | 0.152 | 8.5 g/L | 0.113 | 67 | 353 | 3.32 |
|  | 2 |  | 0.152 | 8.5 g/L | 0.073 | 64 | 320 | 2.68 |
|  | 3 |  | 0.153 | 8.5 g/L | 0.095 | 63 | 325 | 2.92 |
|  | 4 |  | 0.157 | 8.5 g/L | 0.057 | 63 | 313 | 2.63 |
| **Isoprenol TALE IPL300** | 5 | 4 g/L | 0.253 | 8.5 g/L | 0.091 | 74 | 320 | 3.17 |
|  | 6 |  | 0.253 | 8.5 g/L | 0.095 | 70 | 341 | 2.68 |
|  | 7 |  | 0.252 | 8.5 g/L | 0.060 | 64 | 297 | 2.05 |
|  | 8* |  | 0.252 | 7.5 g/L | 0.137 | 29 | 143 | 1.33 |
| **Isoprenol TALE IPL400** | 9 | 4 g/L | 0.300 | 8.5 g/L | 0.085 | 67 | 340 | 2.96 |
|  | 10 |  | 0.300 | 8 g/L | 0.119 | 66 | 326 | 2.80 |
|  | 11 |  | 0.305 | 8 g/L | 0.062 | 51 | 244 | 1.91 |
|  | 12 |  | 0.298 | 8 g/L | 0.061 | 55 | 256 | 1.94 |
| **HCHO TALE KT2440** | 13 | 1 mM | 0.284 | 9 mM | 0.197 | 75 | 387 | 3.56 |
|  | 14 |  | 0.247 | 9 mM | 0.204 | 71 | 363 | 3.35 |
|  | 15 |  | 0.294 | 9 mM | 0.144 | 70 | 362 | 3.41 |
|  | 16 |  | 0.288 | 9 mM | 0.157 | 68 | 346 | 3.11 |

<sup>a</sup>Average growth rate, observed in the three first flasks of each experiment. <sup>b</sup>Average growth rate, observed in the three last flasks of each experiment. \*Has a lower passage than other lineages since the experiment was restarted from an early intermediate stock due to cross-contamination.

**Supplementary Table 4.**
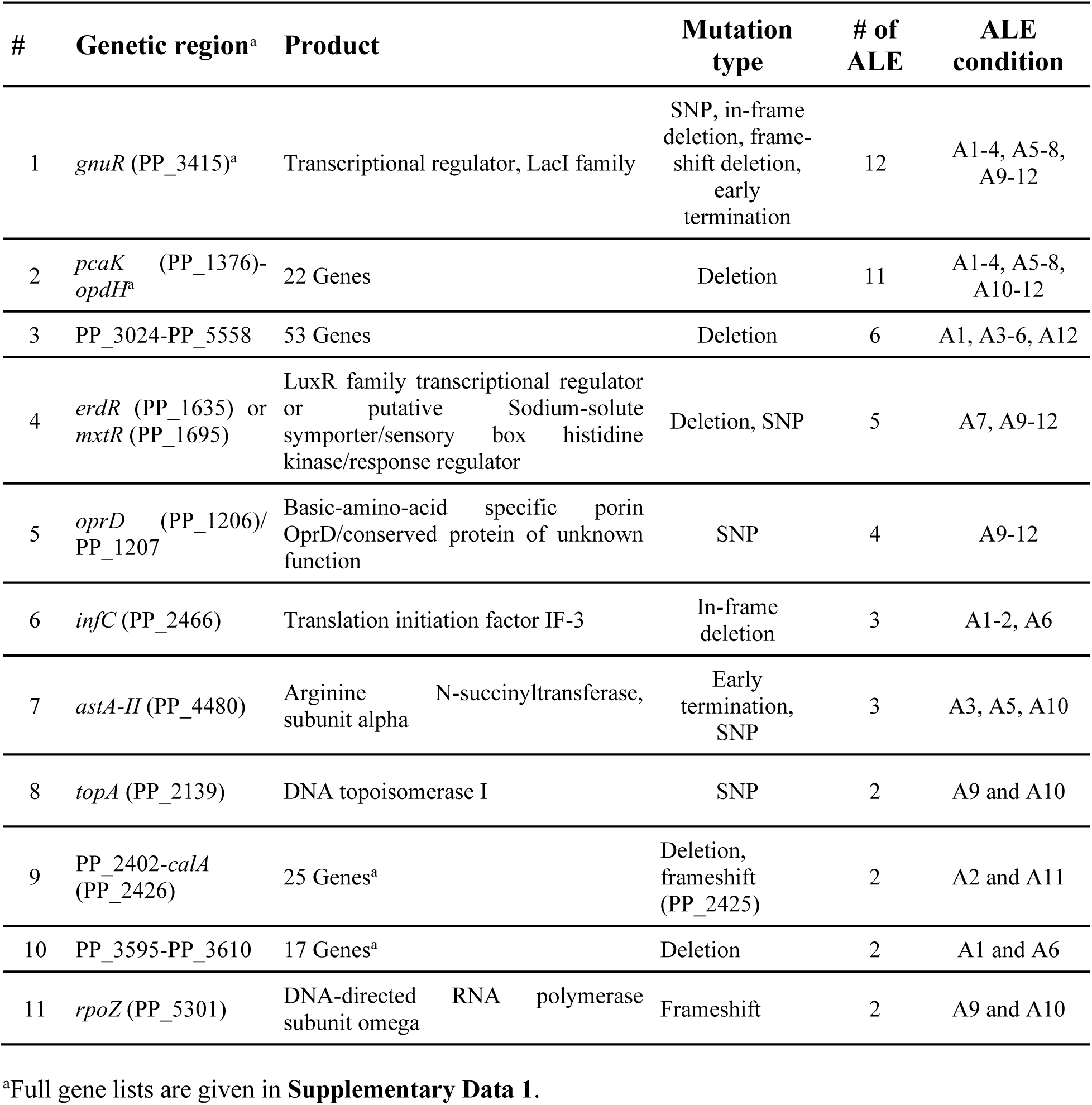
Commonly Mutated Isoprenol Tolerance Specific Regions.

**Supplementary Table 5.** Details of the mutations in the A10_F63_I1 and A12_F53_I1 strains and their corresponding proteomics shifts on glucose minimal medium with respect to IPL400.

| # | Position | Gene | Details | Sequence change | Proteome change |
| --- | --- | --- | --- | --- | --- |
| A10_F63_I1 |  |  |  |  |  |
| 1 | 1,386,816 | *oprD*/PP_1207 | intergenic (-360/-331) | C→T | NS/NS |
| 2 | 1,812,462 | PP_1615/*frmA* | intergenic (-45/-60) | A→G | ND/Up |
| 3 | 1,888,423 | PP_1695 | L617R (CTG→CGG) | A→C | NS |
| 4 | 2,443,213 | *topA* | L784Q (CTG→CAG) | T→A | NS |
| 5 | 3,866,379 | PP_3415 | G167R (GGC→CGC) | C→G | Down |
| 6 | 4,586,057 | PP_4061/PP_4063 | intergenic (+140/+75) | +C | ND/Up |
| 7 | 4,808,180 | PP_4234/*dsbD*-II | intergenic (-289/-265) | 2 bp→CG | NS |
| 8 | 6,049,015 | *rpoZ* | coding (200/264 nt) | +C | NS |
| A12_F53_I1 |  |  |  |  |  |
| 1 | 1,386,816 | *oprD*/PP_1207 | intergenic (-360/-331) | C→T | NS/NS |
| 2 | 1,591,652 | PP_1395 | L81R (CTG→CGG) | A→C | NS |
| 3 | 1,812,462 | PP_1615/*frmA* | intergenic (-45/-60) | A→G | ND/Up |
| 4 | 1,832,579 | PP_1635 | A190T (GCG→ACG) | G→A | NS |
| 5 | 1,851,825 | *relA* | coding (911-922/2241 nt) | Δ12 bp | NS |
| 6 | 2,481,262 | PP_2175 | F29V (TTC→GTC) | A→C | ND |
| 7 | 3,410,136 | PP_3024 | coding (167/1197 nt) | Δ1 bp | Down |
| 8 | 3,866,001 | PP_3415 | P293S (CCA→TCA) | G→A | Down |
| 9 | 3,866,742 | PP_3415 | V46I (GTC→ATC) | C→T |  |
| 10 | 4,586,057 | PP_4061/PP_4063 | intergenic (+140/+75) | +C | ND/Up |

NS: No Significant difference, ND: Not Detected

**Supplementary Table 6.** Summary of regulators enriched in the differentially expressed proteins in the evolved production strain A10_F53_I1_pIY670 vs IPL400.

|  | **pvalue** | **precision** | **recall** | **f1score** | **TP** | **target_**<br>**set_size** | **gene_set**<br>**_size** |
| --- | --- | --- | --- | --- | --- | --- | --- |
| FleQ | 0.0138631806 | 0.07740585774 | 0.1237458194 | 0.09523809524 | 37 | 299 | 478 |
| TurA | 0.01078247117 | 0.06276150628 | 0.1321585903 | 0.08510638298 | 30 | 227 | 478 |
| GacA | 0.1384595471 | 0.04184100418 | 0.1111111111 | 0.06079027356 | 20 | 180 | 478 |
| Crc | 0.04983625899 | 0.02510460251 | 0.1445783133 | 0.04278074866 | 12 | 83 | 478 |
| GlnG | 0.006512587133 | 0.02510460251 | 0.1904761905 | 0.04436229205 | 12 | 63 | 478 |
| AmrZ | 0.6489406466 | 0.0230125523 | 0.07971014493 | 0.03571428571 | 11 | 138 | 478 |
| AlgR | 0.9285961317 | 0.02092050209 | 0.05882352941 | 0.03086419753 | 10 | 170 | 478 |
| Fur | 0.007672164033 | 0.02092050209 | 0.2040816327 | 0.03795066414 | 10 | 49 | 478 |
| PhoB | 0.0008134219476 | 0.01673640167 | 0.32 | 0.03180914513 | 8 | 25 | 478 |
| RoxR | 0.3806535899 | 0.01255230126 | 0.1034482759 | 0.0223880597 | 6 | 58 | 478 |
| PvdS | 0.0003959000084 | 0.01255230126 | 0.4615384615 | 0.02443991853 | 6 | 13 | 478 |
| CbrB | 0.499960924 | 0.01046025105 | 0.09259259259 | 0.01879699248 | 5 | 54 | 478 |
| TurB | 0.4660007998 | 0.01046025105 | 0.09615384615 | 0.01886792453 | 5 | 52 | 478 |
| PcaR | 0.0004314071516 | 0.01046025105 | 0.5555555556 | 0.0205338809 | 5 | 9 | 478 |
| GclR | 0.2201929567 | 0.006276150628 | 0.1578947368 | 0.01207243461 | 3 | 19 | 478 |
| PrpR | 0.01034649773 | 0.006276150628 | 0.5 | 0.01239669421 | 3 | 6 | 478 |
| PydR | 0.005523971961 | 0.006276150628 | 0.6 | 0.01242236025 | 3 | 5 | 478 |
| SoxR | 0.8605034008 | 0.004184100418 | 0.05128205128 | 0.007736943907 | 2 | 39 | 478 |
| PedR1 | 0.8030148711 | 0.004184100418 | 0.05882352941 | 0.0078125 | 2 | 34 | 478 |
| GbdR | 0.7591646544 | 0.004184100418 | 0.06451612903 | 0.007858546169 | 2 | 31 | 478 |
| PsrA | 0.3415729569 | 0.004184100418 | 0.1428571429 | 0.008130081301 | 2 | 14 | 478 |
| FnrA | 0.2427759115 | 0.004184100418 | 0.1818181818 | 0.0081799591 | 2 | 11 | 478 |
| GnuR | 0.2098683974 | 0.004184100418 | 0.2 | 0.008196721311 | 2 | 10 | 478 |
| AcoR | 0.1158454094 | 0.004184100418 | 0.2857142857 | 0.00824742268 | 2 | 7 | 478 |
| LldR | 0.03931609895 | 0.004184100418 | 0.5 | 0.008298755187 | 2 | 4 | 478 |
| LexA-I | 0.8489121889 | 0.002092050209 | 0.04761904762 | 0.004008016032 | 1 | 21 | 478 |
| PplR1 | 0.7833207891 | 0.002092050209 | 0.05882352941 | 0.00404040404 | 1 | 17 | 478 |
| PP_3603 | 0.7405421275 | 0.002092050209 | 0.06666666667 | 0.004056795132 | 1 | 15 | 478 |
| PP_4759 | 0.7405421275 | 0.002092050209 | 0.06666666667 | 0.004056795132 | 1 | 15 | 478 |
| RpoS | 0.7160900646 | 0.002092050209 | 0.07142857143 | 0.00406504065 | 1 | 14 | 478 |
| HexR | 0.5547143094 | 0.002092050209 | 0.1111111111 | 0.004106776181 | 1 | 9 | 478 |
| GltR-II | 0.5127988781 | 0.002092050209 | 0.125 | 0.004115226337 | 1 | 8 | 478 |

|  | **pvalue** | **precision** | **recall** | **f1score** | **TP** | **target_**<br>**set_size** | **gene_set**<br>**size** |
| --- | --- | --- | --- | --- | --- | --- | --- |
| PtxS | 0.5127988781 | 0.002092050209 | 0.125 | 0.004115226337 | 1 | 8 | 478 |
| RutR-I | 0.4669469119 | 0.002092050209 | 0.1428571429 | 0.00412371134 | 1 | 7 | 478 |
| RbsR | 0.4669469119 | 0.002092050209 | 0.1428571429 | 0.00412371134 | 1 | 7 | 478 |
| FruR | 0.3619234664 | 0.002092050209 | 0.2 | 0.004140786749 | 1 | 5 | 478 |
| DctD-II | 0.3619234664 | 0.002092050209 | 0.2 | 0.004140786749 | 1 | 5 | 478 |
| GlcC | 0.3619234664 | 0.002092050209 | 0.2 | 0.004140786749 | 1 | 5 | 478 |
| NrdR | 0.3019076099 | 0.002092050209 | 0.25 | 0.004149377593 | 1 | 4 | 478 |
| CueR | 0.3019076099 | 0.002092050209 | 0.25 | 0.004149377593 | 1 | 4 | 478 |
| GcsR | 0.3019076099 | 0.002092050209 | 0.25 | 0.004149377593 | 1 | 4 | 478 |
| GlpR | 0.3019076099 | 0.002092050209 | 0.25 | 0.004149377593 | 1 | 4 | 478 |
| FleN | 0.3019076099 | 0.002092050209 | 0.25 | 0.004149377593 | 1 | 4 | 478 |
| OxyR | 0.3019076099 | 0.002092050209 | 0.25 | 0.004149377593 | 1 | 4 | 478 |
| SdaR | 0.2362597321 | 0.002092050209 | 0.3333333333 | 0.004158004158 | 1 | 3 | 478 |
| DctD-III | 0.08590941769 | 0.002092050209 | 1 | 0.004175365344 | 1 | 1 | 478 |

Data compared against the transcriptional regulation network database reported in Lim et al., 2022 (doi:10.1016/j.ymben.2022.04.004)

